# Indirect CRISPR screening with photoconversion revealed key factors of drug resistance with cell–cell interactions

**DOI:** 10.1101/2022.07.15.500173

**Authors:** Keisuke Sugita, Iichiroh Onishi, Ran Nakayama, Sachiko Ishibashi, Masumi Ikeda, Miori Inoue, Rina Narita, Shiori Oshima, Kaho Shimizu, Shinichiro Saito, Shingo Sato, Branden S. Moriarity, Kouhei Yamamoto, David A. Largaespada, Masanobu Kitagawa, Morito Kurata

## Abstract

Comprehensive screenings to clarify indirect cell–cell interactions, such as those in the tumor microenvironment, especially comprehensive assessments of supporting cells’ effects, are challenging. Therefore, in this study, indirect CRISPR screening for drug resistance with cell–cell interactions was invented. The photoconvertible fluorescent protein Dendra2 was inducted to determine the drug resistance responsible factors of supporting cells with CRISPR screenings. Random mutated supporting cells co-cultured with leukemic cells induced drug resistance with cell– cell interactions. Supporting cells responsible for drug resistance were isolated with green-to-red photoconversion, and 39 candidate genes were identified. Knocking out *C9orf89*, *MAGI2*, *MLPH*, or *RHBDD2* in supporting cells reduced the ratio of apoptosis of cancer cells. In addition, the low expression of RHBDD2 in supporting cells, specifically fibroblasts, of clinical pancreatic cancer showed a shortened prognosis, and a negative correlation with CXCL12 was observed. Indirect CRISPR screening was established to isolate the responsible elements of cell–cell interactions. This screening method could reveal new mechanisms in all kinds of cell–cell interactions by revealing live phenotype-inducible cells, and it could be a new platform for discovering new targets of drugs for conventional chemotherapies.

## Introduction

Many oncogenes and tumor suppressor genes have been discovered, and anticancer drugs, including molecularly targeted drugs, are now widely used. However, cancer is still resistant to treatment, and drug resistance remains a major concern in discovering cures (Chen & Song, 2019) (Vasan et al., 2019) (Kanzaki & Pietras, 2020). Therefore, the elucidation of drug resistance mechanisms is an important challenge to overcome. Regarding drug resistance systems, drug inactivation, drug target alteration, drug efflux, DNA damage repair, cell death inhibition, and the epithelial-mesenchymal transition are well-known and well-studied (Housman et al., 2014). In addition to these mechanisms, the tumor microenvironment (TME), the peritumoral region composed of cancer-associated fibroblasts (CAFs), the extracellular matrix (ECM), immune cells, and vasculature have been the focus of research in recent years, as they co-evolve during malignant progression and contribute to cancer development, progression, and drug resistance (Vasan et al., 2019) (Kanzaki & Pietras, 2020) (Hosein et al., 2020).

The microenvironment was originally studied as part of the bone marrow as the bone marrow niche that maintains and regulates hematopoietic stem cells, and TME is also involved in the drug resistance of myeloma in the bone marrow, aside from solid tumors (Morrison & Scadden, 2014) (Meads et al., 2008) (Danziger et al., 2020). CAFs are also known to be stromal cells of the TME that have distinctive properties compared to fibroblasts derived from normal tissue and have been implicated in tumor growth, local invasion, distant metastasis, ECM remodeling, angiogenesis, and drug resistance (Chen & Song, 2019) (Orimo et al., 2005) (Kalluri, 2016) (Sahai et al., 2020) (Wu et al., 2021). Because of the wide variety of drug resistance mechanisms associated with cell–cell interactions in the TME, comprehensive analyses of these mechanisms are crucial for overcoming drug resistance.

According to global cancer statistics (GLOBOCAN 2020), pancreatic cancer has a poor prognosis, with the number of related deaths (466,003) being almost equal to the number of patients (495,773), and it is the seventh leading cause of cancer deaths in both men and women (Sung et al., 2021). The 5-year survival rate is approximately 3–10%, and 80–85% of patients are ineligible for surgical resection at the time of diagnosis due to either the state of the locally advanced disease or distant metastases (Siegel et al., 2018) (Peng et al., 2019) (Mizrahi et al., 2020) (Singhi et al., 2019). Pancreatic ductal adenocarcinoma (PDAC) is characterized by abundant desmoplastic stroma, and the TME in this stroma promotes drug resistance and a worse prognosis in PDAC (Hosein et al., 2020) (Ogawa et al., 2021) (Grünwald et al., 2021).

In this study, an experimental model using CRISPR screening was created to more comprehensively elucidate the mechanism behind drug resistance induction by peritumoral supporting cells, whereas previous reports on TME or CAFs have mainly focused on TME tissue derived from “poor prognostic cancers” (Orimo et al., 2005) (Grünwald et al., 2021) (Su et al., 2018) (Uchihara et al., 2020).

In conventional CRISPR screening, the candidate genes for drug resistance are identified by detecting the increasing/decreasing number of tumor cells with guide RNAs (gRNAs) that induce random mutations under screening conditions, such as drug exposure (Kurata et al., 2018). However, in cases of drug resistance by cell–cell interactions from the surrounding environment, such as the TME, the identification of the responsible candidate genes for drug resistance with cell–cell interactions is difficult. This is because there is no selective increase in the number of surrounding cells, even if random mutations with gRNAs are induced in the surrounding cells (Figure 1A). In this study, a new CRISPR screening system, indirect CRISPR screening, using the photoconvertible fluorescent protein Dendra2 was established to identify new responsible molecules in drug resistance induced by peritumoral cell–cell interactions.

**Figure 1.**
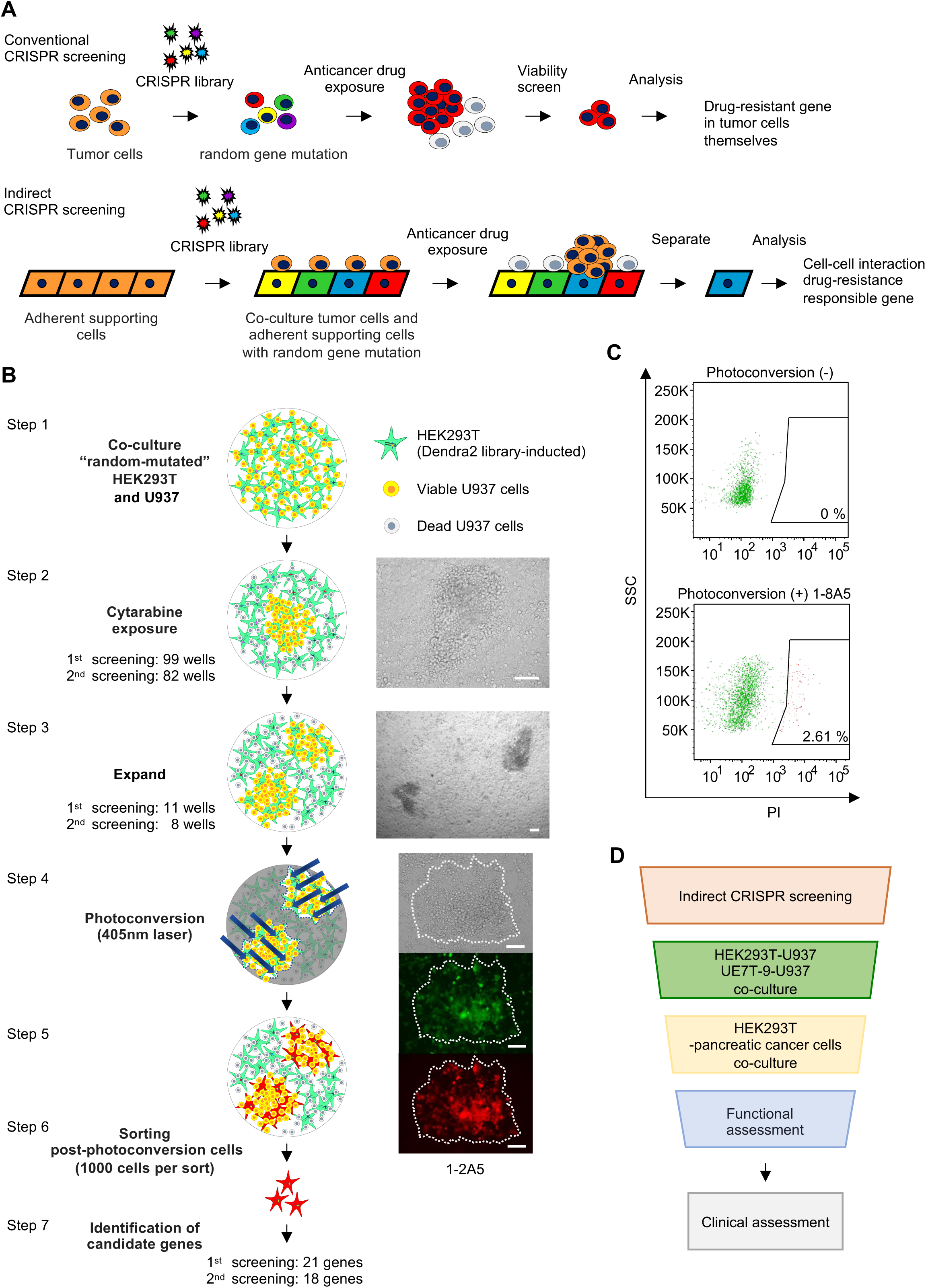
Indirect CRISPR screening system with photoconversion. (**A**) The experimental model. In conventional CRISPR screening, the candidate genes for drug resistance are identified by detecting the increasing/decreasing number of tumor cells with gRNAs inducing random mutations under screening conditions (upper tier). Meanwhile, in indirect CRISPR screening, random mutations are inducted into the adherent supporting cells and co-cultured with the tumor cells to create so-called “microenvironments.” The responsible supporting cells are then separated and analyzed for drug resistance with cell–cell interactions (lower tier). (**B**) Indirect CRISPR screening system with Dendra2. HEK293T and U937 cells were inducted as cell–cell interaction systems to mimic the microenvironment. U937 was co-cultured with HEK293T inducted with Dendra2 and the CRISPR library (Step 1). Under cytarabine exposure, most U937 cells were killed, and only some U937 cells close to HEK293T survived and proliferated (Step 2). The supporting cells that induced drug resistance in tumor cells were expanded (Step 3) and then identified by laser scanning microscopy. They then underwent photoconversion with the 405 nm laser (Steps 4 and 5). After photoconversion, the red-fluorescing HEK293T cells were sorted on FACS (Step 6). Sorted cells were analyzed to identify the genes responsible for drug resistance induced by cell–cell interactions (Step 7). Microscopic images are representative images corresponding to each schema (bright-field image and laser scanning images; green: 473 nm, red: 559 nm). (**C**) FACS sorted supporting cells without (upper tier) or with (lower tier) photoconversion using the PI filter. Photoconverted cells were sorted by 1,000 cells per well. (**D**) Pipeline of validation experiment. Scale bar, 50 μm.

## Results

### Indirect CRISPR screening with photoconversion

In our new experimental model, random mutations were inducted into supporting cells and co-cultured with the tumor cells to mimic the microenvironments of drug resistance and to identify the responsible cells using photoconversion and isolation. To identify and isolate cells that support drug resistance using the CRISPR library, we utilized a photoconversion protein, Dendra2, which can irreversibly convert its fluorescence wavelength from green to red with illumination from ultraviolet (UV) light (405 nm laser) (Figure 1—figure supplement 1) (Gurskaya et al., 2006).

HEK293T and U937 leukemia cells, which were easy to handle technically, were inducted as cell– cell interaction systems to mimic a microenvironment. Dendra2-inducted HEK293T cells, with the CRISPR knockout library also inducted, were co-cultured with U937 cells (Figure 1B, Step 1). U937 is relatively more sensitive to cytarabine than HEK293T (Figure 2—figure supplement 1A, B), and co-culture screening was performed with 3 μM cytarabine. Most U937 cells were killed, but a small number of U937 in close proximity to HEK293T survived and proliferated in a mulberry-like manner (Figure 1B, Step 2). The CRISPR library induction to HEK293T cells and screening were performed independently twice. The screenings were performed with 7.2 × 10^6^ cells of HEK293T and 1.44 × 10^7^ cells of U937 in 15 96-well plates per each screening. The survival of U937 colonies in 96-well plates was identified in 99 wells from the first screening and in 82 wells from the second screening. To obtain sufficient supporting cells for analysis and to confirm their ability to induce drug resistance reproducibly, all cells in the viable U937 colony-positive wells were transferred and expanded on a large scale. The colonies of positive wells were re-seeded on 6-well plates at 11 wells from the first screening and 8 wells from the second screening (Figure 1B, Step 3). All supporting cells close to the viable U937 colonies under cytarabine exposure observed in the 19 wells were applied to photoconversion. The supporting cells that induced drug resistance near viable U937 colonies were identified by differential interference images from laser scanning microscopy, and photoconversion was performed with the 405 nm laser (Figure 1B, Step 4). After photoconversion, the red form of Dendra2 was confirmed with laser scanning microscopy (559 nm laser) (Figure 1B, Step 5) and the PI filter of FACS. Then, 1,000 HEK293T cells with the red form of Dendra2 were sorted from each well by FACS (Figure 1B, Step 6 and 1C).

**Figure 2.**
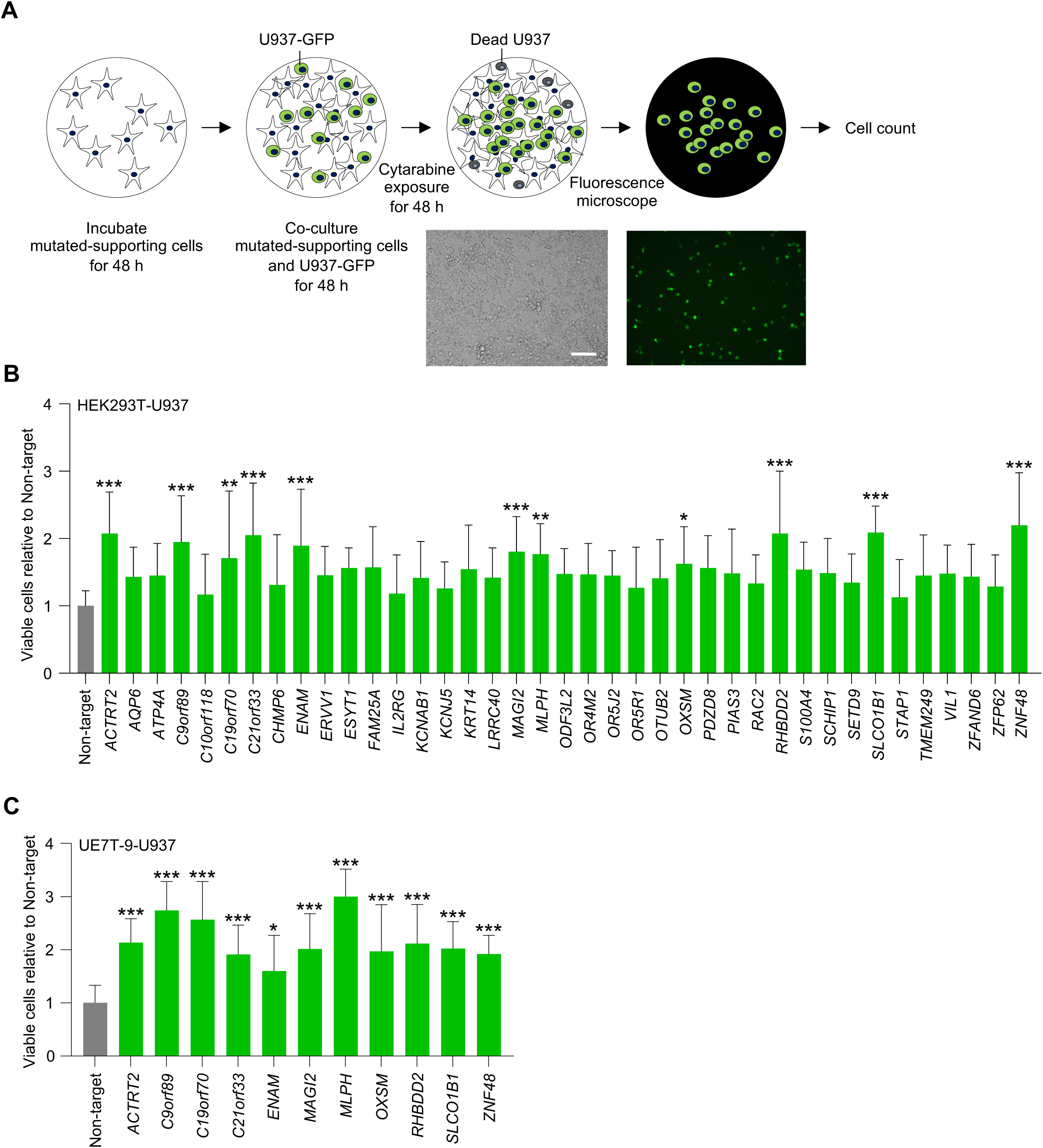
**Validation of drug resistance with cell–cell interactions in HEK293T-U937 and UE7T**-**9-U937 model.** (**A**) Experimental scheme for the detection of living U937 cells in co-culture experiments. (**B-C**) The cell count of viable U937 cells co-cultured with knockout mutant-supporting cells (HEK293T (B) and UE7T-9 (C)) treated with 5 μM of cytarabine for 48 h. Three fields (20x objective field) were randomly captured for each well, and the numbers of viable U937 cells were counted. The x-axis represents the knockout candidate genes of the supporting cells. The y-axis represents the number of viable cells relative to that of co-cultured cells with the control (non-targeted gRNA) of supporting cells. All experiments were performed in biological triplicate in each of the two independent experiments. Data are represented as mean ± SD. Statistical significance values were calculated by performing one-way ANOVA using Dunnett’s test (B-C). \**p* < 0.05; \*\**p* < 0.01 and \*\*\**p* < 0.001. Scale bar, 50 μm. Figure 2**—source data 1.** Values for the graph in Figure 2B. Figure 2**—source data 2.** Values for the graph in Figure 2C.

Collected mutated HEK293T cells were then expanded, and mutations were analyzed, after which 39 candidate genes were obtained (Figure 1B, Step 7 and Table 1). Of the 39 genes, 21 were from the first screening, and 18 were from the second screening. These 39 candidate genes were validated to determine whether they could reproducibly induce drug resistance and associated mechanisms (Figure 1D).

**Table 1.**
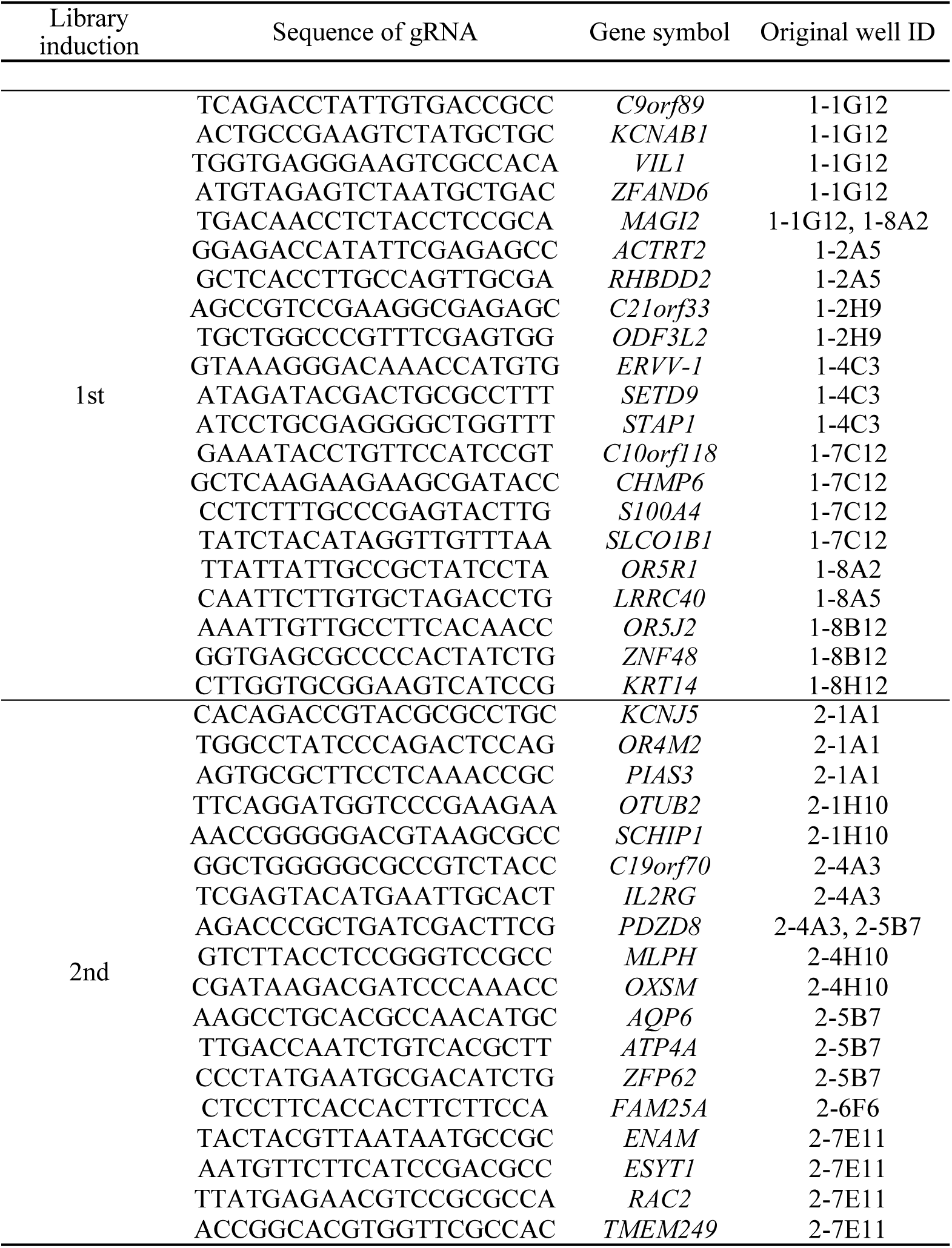
Guide RNAs in HEK293T clones inducing drug resistance with cell–cell interactions

### Validation of leukemic drug resistance induced by supporting cells

To validate the drug resistance functions of the 39 candidate genes in cell–cell interactions, each candidate gene in HEK293T was knocked out, and GFP-positive U937 cells were co-cultured to estimate the number of surviving tumor cells under cytarabine exposure using immunofluorescent microscopy (Figure 2A). gRNA-inducted HEK293T and GFP-positive U937 were co-cultured and exposed to 5 µM of cytarabine (Figure 2—figure supplement 1A, B) in 24-well plates for 48 h. The fluorescent images were evaluated, and the number of viable GFP-positive U937 cells was assessed (Figure 2B and Figure 2—figure supplement 2). The results showed that the surviving number of viable U937 cells was significantly increased when co-cultured with HEK293T gRNA inducted for the following 11 candidate genes: *ACTRT2*, *C9orf89*, *C19orf70*, *C21orf33*, *ENAM*, *MAGI2*, *MLPH, OXSM*, *RHBDD2*, *SLCO1B1*, and *ZNF48*.

To confirm the universality (i.e., not specific in HEK293T cells) of the gene functions of these 11 candidates in cell–cell interactions, the human bone marrow-derived mesenchymal stem cell (UE7T-9) was used instead of HEK293T, which was used as an experimental model for cell–cell interactions. The same experiment was conducted to verify whether the function of drug resistance could be induced in the co-culture experiment with UE7T-9 and U937 (Figure 2C, Figure 2—figure supplement 1B, C, and Figure 2—figure supplement 3). The results showed that the number of surviving GFP-positive U937 cells increased significantly with UE7T-9 by knocking out the 11 candidate genes of *ACTRT2, C9orf89, C19orf70, C21orf33, ENAM, MAGI2, MLPH, OXSM, RHBDD2, SLCO1B1,* and *ZNF48,* respectively. Therefore, the above 11 candidate genes could induce universal drug resistance with cell–cell interactions and were explored further.

### Application to pancreatic models of drug resistance with cell–cell interactions

Pancreatic cancer is associated with abundant stroma, and this peritumoral stroma is associated with drug resistance (Ogawa et al., 2021). Additionally, cytarabine and gemcitabine are both pyrimidine analogues and have similar chemical structures (Derissen & Beijnen, 2020). Therefore, candidate genes that induced cytarabine resistance were also evaluated to determine whether they induced gemcitabine resistance. The candidate genes that induced drug resistance in U937 cells in co-culture with HEK293T or UE7T-9 cells were applied to assess whether they universally induce drug resistance with cell–cell interactions in other cancers. Invasive pancreatic ductal carcinoma cell lines MIA PaCa-2 and SUIT-2 with HEK293T and the anticancer drug gemcitabine were conducted in co-culture experiments.

As the sensitivity among HEK293T, MIA PaCa-2, and SUIT-2 to gemcitabine was comparable, *Deoxycytidine kinase* (*DCK*) knockout HEK293T was generated. DCK deficiency is known to be a major factor in gemcitabine resistance *in vitro* and *in vivo* due to its central role in gemcitabine metabolism (Binenbaum et al., 2015). Cloning of knockout HEK293T cells was performed using limiting dilution methods, and CRISPR knockout clones (CKO) were established (Figure 3—figure supplement 1A). IC50 against gemcitabine was remarkably increased in HEK293T *DCK*-CKO compared to MIA PaCa-2 or SUIT-2 (HEK293T: 1.46 × 10^-3^ μM, HEK293T *DCK*-CKO: 23.8 μM, MIA PaCa-2: 4.10 × 10^-2^ μM, SUIT-2: 8.33 × 10^-3^ μM, Figure 3—figure supplement 1B-E). To differentiate between HEK293T and pancreatic cancer cell lines under fluorescence microscopy, only pancreatic cancer cells were transfected with GFP for co-culture and gemcitabine exposure experiments (Figure 3—figure supplement 2).

**Figure 3.**
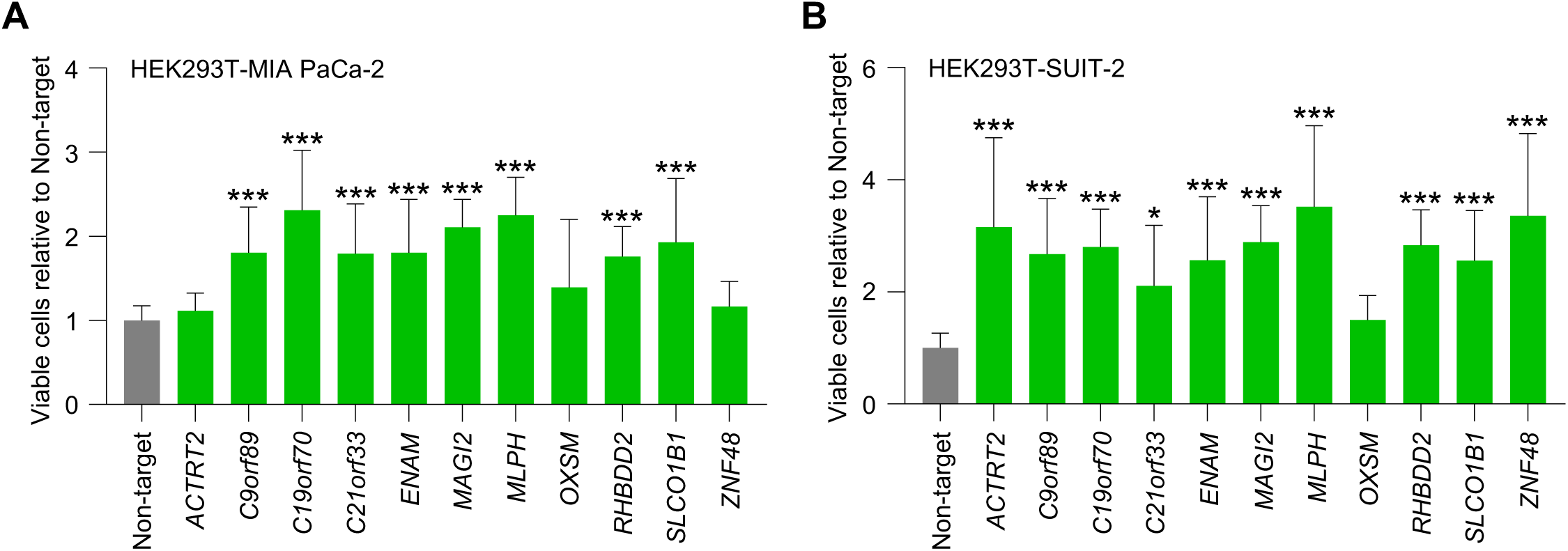
Validation of drug resistance with cell–cell interactions in HEK293T-pancreatic cancer cells model. (**A-B**) The cell counts of viable pancreatic cancer cells (MIA PaCa-2 (A) and SUIT-2 (B)) co-cultured with knockout mutant-HEK293T treated with gemcitabine (10 μM (A) or 3 μM (B)) for 48 h. Three fields (20x objective field) were randomly captured for each well, and the numbers of viable pancreatic cancer cells were counted. The x-axis represents the knockout candidate genes of HEK293T. The y-axis represents the number of viable cells relative to that co-cultured with the control (non-targeted gRNA) of HEK293T. All experiments were performed in biological triplicate in each of the two independent experiments. Data are represented as mean ± SD. Statistical significance values were calculated by performing one-way ANOVA using Dunnett’s test (A-B). \**p* < 0.05 and \*\*\**p* < 0.001. Figure 3**—source data 1.** Values for the graph in Figure 3A. Figure 3**—source data 2.** Values for the graph in Figure 3B.

The number of surviving GFP-positive pancreatic cancer cells under gemcitabine exposure increased significantly universally in MIA PaCa-2 (Figure 3A and Figure 3—figure supplement 3A) and SUIT-2 (Figure 3B and Figure 3—figure supplement 3B) when co-cultured with HEK293T, which was knocked out for eight candidate genes: *C9orf89, C19orf70, C21orf33, ENAM, MAGI2, MLPH, RHBDD2,* and *SLCO1B1*. The knockout of these eight genes in the supporting cells universally induced the survival of leukemia cells under cytarabine and pancreatic cancer cells under gemcitabine.

### Addressing biologically responsible factors in drug resistance with cell–cell interactions

To assess the universal and obvious effects of drug resistance with cell–cell interactions, highly expressed candidate genes in HEK293T, specifically *C9orf89, C19orf70, C21orf33, MAGI2, MLPH*, and *RHBDD2* (Figure 4—figure supplement 1), were focused on and applied to the next experiments.

**Figure 4.**
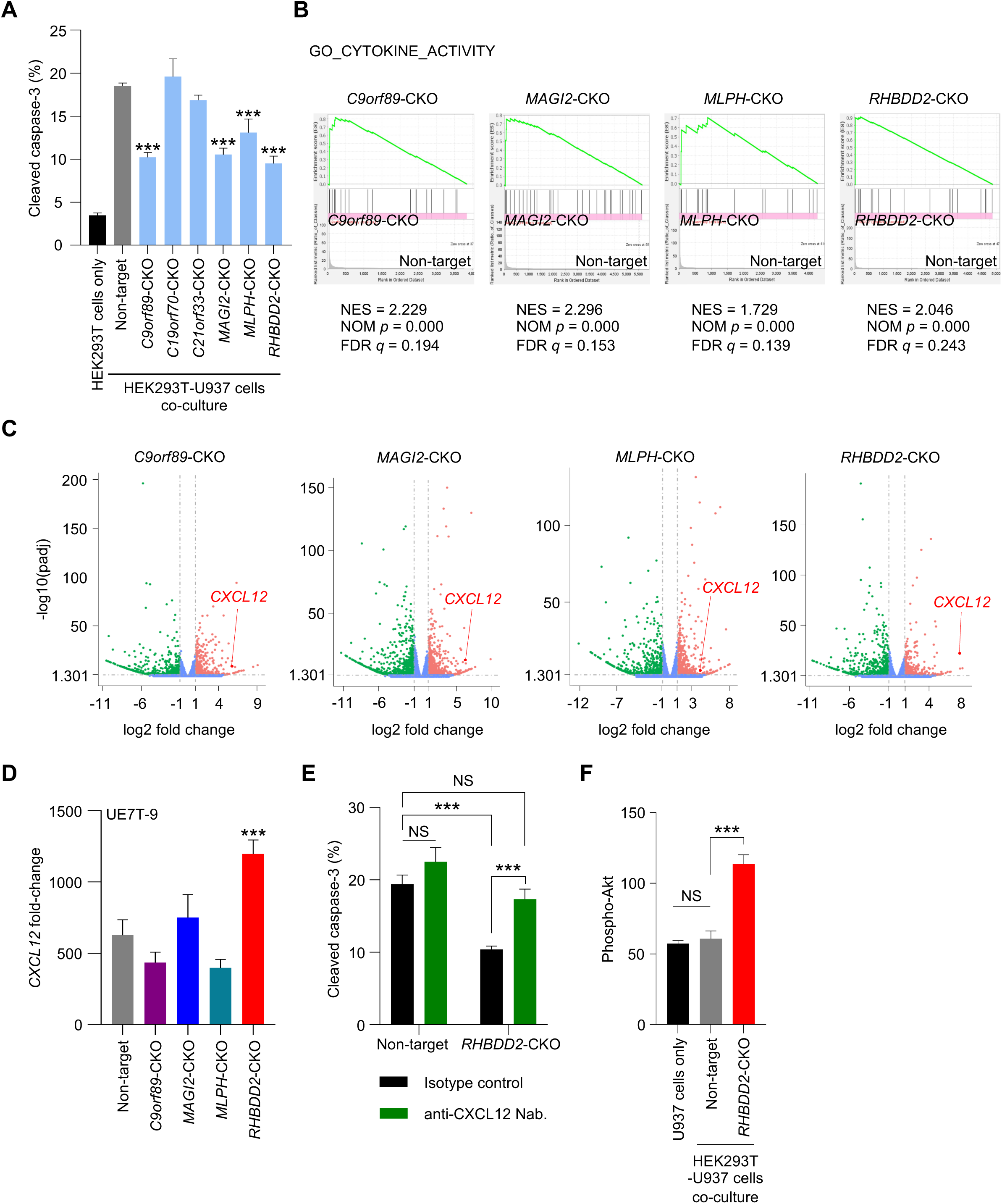
Addressing biologically responsible factors in drug resistance with cell–cell interactions with RNA-seq and the regulation of *CXCL12* in *RHBDD2*-CKO cells. (**A**) Ratio of apoptotic cells in the co-culture system with *C9orf89*-CKO, *MAGI2*-CKO, *MLPH*-CKO, and *RHBDD2*-CKO HEK293T under cytarabine exposure for 48 h. (**B**) GSEA of HEK293T-CKO clones vs. HEK293T control showing enrichment of gene sets involved in cytokine activity. (**C**) Volcano plot of HEK293T-CKO clones vs. HEK293T control showing the up-regulation of *CXCL12 in C9orf89*-CKO, *MAGI2*-CKO, *MLPH*-CKO, and *RHBDD2*-CKO HEK293T. (**D**) Different cell types of UE7T-9 showing elevated *CXCL12* expression in *RHBDD2*-CKO UE7T-9. (**E**) In the co-culture experiment of *RHBDD2*-CKO HEK293T and U937, pre-treatment with CXCL12-neutralizing antibodies increased cleaved caspase-3-positive cells under cytarabine exposure for 48 h, which was similar to that observed when the cells were co-cultured with control HEK293T. (**F**) Evaluation of phospho-Akt, downstream of CXCL12, in GFP-positive U937 under mono- or co-culture conditions. Phospho-Akt was increased in U937 co-cultured with *RHBDD2*-CKO HEK293T for 48 h. Each experiment was performed with biological triplication in three independent repeats. Data are represented as mean ± SD. Statistical significance values were calculated by performing one-way ANOVA with Dunnett’s test (A, D, F) and one-way ANOVA with Bonferroni’s test (E). \*\*\**p* < 0.001 and NS: non-significant. Figure 4**—source data 1.** Values for the graph in Figure 4A. Figure 4**—source data 2.** Values for the graph in Figure 4D. Figure 4**—source data 3.** Values for the graph in Figure 4E. Figure 4**—source data 4.** Values for the graph in Figure 4F.

CKOs of HEK293T were established for six candidates (Figure 4—figure supplement 2A, B) and co-cultured with U937, and cytarabine-induced apoptosis was assessed using cleaved caspase-3 (Figure 4A). In HEK293T alone under cytarabine, very few cleaved caspase-3-positive cells were co-cultured with HEK293T cells of *C9orf89*-CKO, *MAGI2*-CKO, *MLPH*-CKO, or *RHBDD2*-CKO compared to the control.

To investigate the biological features of *C9orf89*, *MAGI2*, *MLPH*, and *RHBDD2*, RNA sequences were performed on HEK293T cells of *C9orf89*-CKO, *MAGI2*-CKO, *MLPH*-CKO, and *RHBDD2*-CKO by next-generation sequencers. In gene set enrichment analysis (GSEA), 66 categories were universally up-regulated in HEK293T cells of *C9orf89*-CKO (Supplementary File 1), *MAGI2*-CKO (Supplementary File 2), *MLPH*-CKO (Supplementary File 3), and *RHBDD2*-CKO (Supplementary File 4). In these 66 categories (Supplementary File 5), two categories that had been suggested to be related in drug resistance induced in the microenvironment based on previous studies were “cytokine activity” (Figure 4B) and “cell adhesion mediated by integrin” (Chen & Song, 2019) (Kalluri, 2016) (Wu et al., 2021). In these two categories, *CXCL12* in “cytokine activity” was the most commonly enriched in the four HEK293T-CKO clones based on the rank metric score in GSEA (Supplementary File 6 and Supplementary File 7). *CXCL12* was commonly up-regulated in HEK293T cells of *C9orf89*-CKO, *MAGI2*-CKO, *MLPH*-CKO, and *RHBDD2*-CKO (Figure 4C), therefore was focused on in subsequent experiments. To confirm the expression of *CXCL12* in CKO clones, qPCR was performed. *CXCL12* expression was significantly up-regulated in HEK293T cells of *C9orf89*-CKO, *MAGI2*-CKO, and *RHBDD2*-CKO (Figure 4—figure supplement 3). To confirm the cell–cell universality of elevated *CXCL12* expression, *CXCL12* expression was examined by qPCR in UE7T-9 cells with CKO mutations of these four candidates (Figure 4—figure supplement 2C, D). UE7T-9 *RHBDD2*-CKO showed elevated *CXCL12* expression, while UE7T-9 cells of *C9orf89*-CKO, *MAGI2*-CKO, and *MLPH*-CKO showed no up-regulation of *CXCL12* (Figure 4D). These results confirm that *RHBDD2*-CKO might up-regulate *CXCL12* expression in any type of stromal cell.

### *RHBDD2* regulates CXCL12 expression in supporting cells and CXCL12 induces anticancer drug resistance via the PI3k-Akt-mTOR pathway

To verify that the cytarabine resistance of U937 is induced by the secretion of *RHBDD2*-CKO HEK293T, U937 was cultured with supernatant from *RHBDD2*-CKO HEK293T and exposed to 250 nM of cytarabine. The viability of U937 cultured with the *RHBDD2*-CKO culture supernatant was slightly but significantly increased under cytarabine exposure (Figure 4—figure supplement 4).

To verify that cytarabine resistance of U937 induced by the co-culture with *RHBDD2*-CKO HEK293T is mediated by CXCL12, cytarabine exposure experiments were also performed after pre-treatment with the CXCL12-neutralizing antibody. The results showed that cytarabine exposure following the treatment with CXCL12-neutralizing antibodies increased cleaved caspase-3 positive cells under cytarabine exposure, which was similar to that observed when the cells were co-cultured with control HEK293T (Figure 4E).

A known pathway for the induction of CXCL12-mediated anti-cancer drug resistance is the activation of the PI3K-Akt-mTOR system in tumor cells (Wu et al., 2021). Therefore, whether elevated CXCL12 in *RHBDD2*-CKO HEK293T results in the activation of the PI3K-Akt-mTOR system in U937 was tested. Phosphorylation of Akt in U937 cells under co-culture with HEK293T cells was evaluated. Increased phosphorylation of Akt in U937 (labeled with GFP to differentiate from HEK293T) cells co-cultured with HEK293T was assessed by flow cytometry. The results showed that phosphorylated-Akt (phospho-Akt) was increased in U937 co-cultured with *RHBDD2*-CKO HEK293T (Figure 4F).

### Analysis of the expression of cell–cell interaction factors in clinical pancreatic cancers

To evaluate whether four important candidate genes in stromal cells, *C9orf89*, *MAGI2*, *MLPH*, and *RHBDD2*, are associated with prognosis, immunohistochemical staining of C9orf89, MAGI2, MLPH, or RHBDD2 in pancreatic ductal carcinoma surgical resection samples without neoadjuvant chemotherapy (n = 60) was conducted (Figure 5–Source Data 1). Immunostainability of fibroblasts surrounding pancreatic carcinoma cells was evaluated. The stainabilities of the cells in the peritumoral stroma were scored as follows: not stained in fibroblasts was (-), positive images in only a few fibroblasts or weakly positive images were (1+), and strongly positive on most of the fibroblasts was (2+). Based on scoring, (-) and (1+) were classified as negative groups, and (2+) was the positive group (Figure 5A and Figure 5—figure supplement 1). Overall survival (OS) was significantly shortened in the negative group compared to the positive group that expressed MAGI2 and RHBDD2 (MAGI2: *p* = 0.014, RHBDD2: *p* < 0.001) (Figure 5B and Table 2). However, for the MLPH and C9orf89 groups, no significant correlation between immunostainability and OS could be identified (MLPH: *p* = 0.514, C9orf89: *p* = 0.089). Six parameters, including age, pT category, tumor size, UICC stage, MAGI2, and RHBDD2, were tested by multivariate analysis using the Cox proportional hazard model (Table 2). The results showed that the pT category was 3–4 (vs. 1–2, hazard ratio: 8.665, *p* = 0.013), UICC stage was III–IV (vs. I–IIB, hazard ratio: 3.051, *p* = 0.003), and the RHBDD2-negative group (vs. RHBDD2-positive group, hazard ratio: 13.590, *p* < 0.001) independently predicted shorter OS.

**Figure 5.**
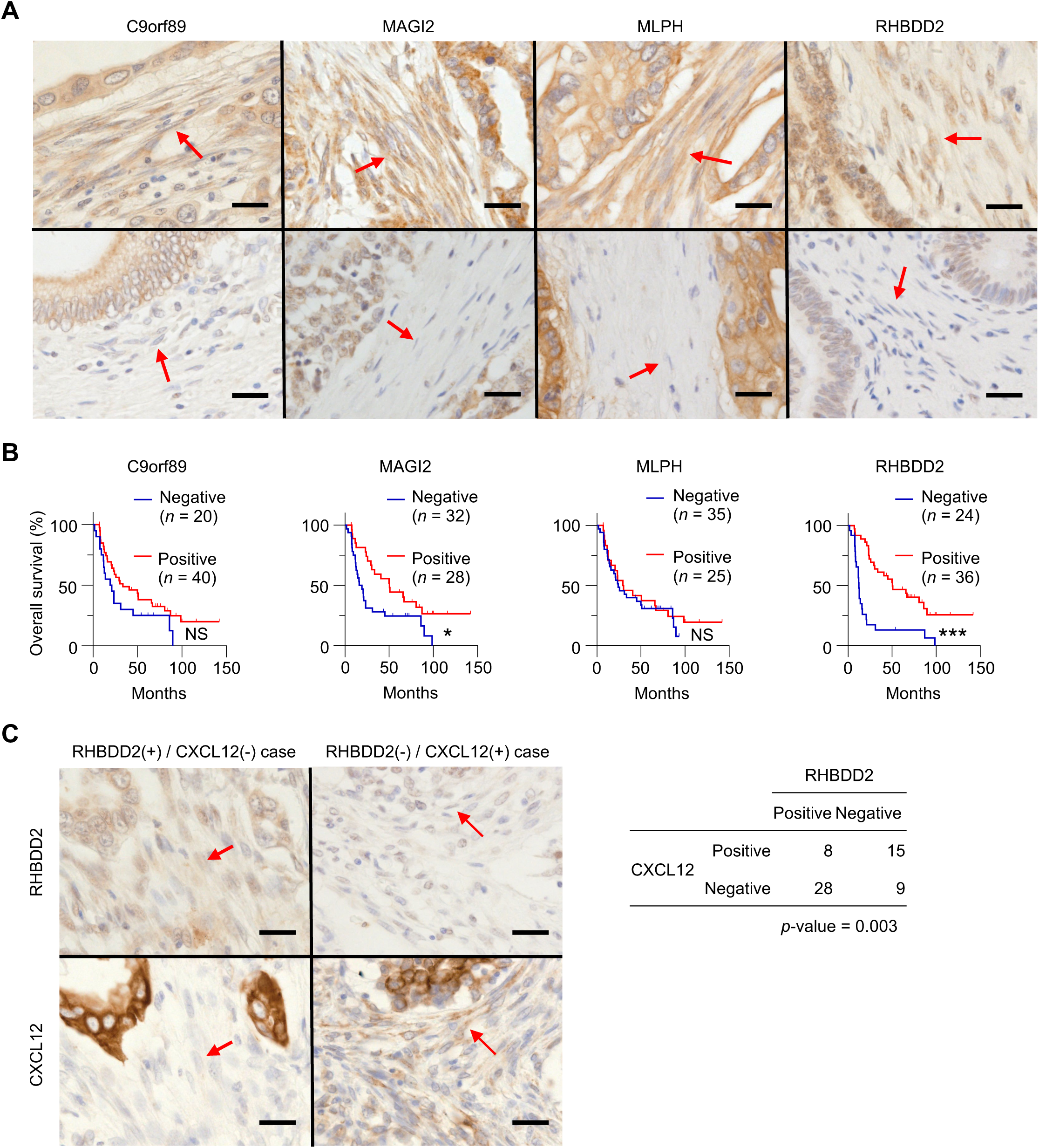
The expression of C9orf89, MAGI2, MLPH, and RHBDD2 in fibroblasts of human pancreatic cancers and their significance as prognostic factors. (**A**) Immunohistochemical staining of C9orf89, MAGI2, MLPH, or RHBDD2 in pancreatic ductal carcinoma surgical resection samples without neoadjuvant chemotherapy (n = 60). The immunostainability of fibroblasts surrounding pancreatic carcinoma cells was evaluated. Representative positive cases (upper tier) and negative cases (lower tier). (**B**) Kaplan–Meier survival curves of pancreatic ductal carcinoma patients with negative and positive groups for C9orf89, MAGI2, MLPH, or RHBDD2 in fibroblasts surrounding carcinoma cells (n = 60). The y-axis signifies elapsed months from the date of diagnosis. (**C**) Association between RHBDD2 and CXCL12 expression in fibroblasts surrounding carcinoma cells. The inverse correlation between RHBDD2 and CXCL12 was found (right table). Left column: a representative case of RHBDD2 (+) / CXCL12 (-); Right column: a representative case of RHBDD2 (+) / CXCL12 (-). Statistical significance values were calculated by performing log-rank tests (B) and Fisher’s exact test (C). \**p* < 0.05; \*\*\**p* < 0.001; and NS: non-significant. Red arrows indicate fibroblasts. Scale bar, 25 μm. Figure 5**—source data 1.** Summary of patient clinical records and immunostainability of peritumoral fibroblasts (n = 60).

**Table 2.**
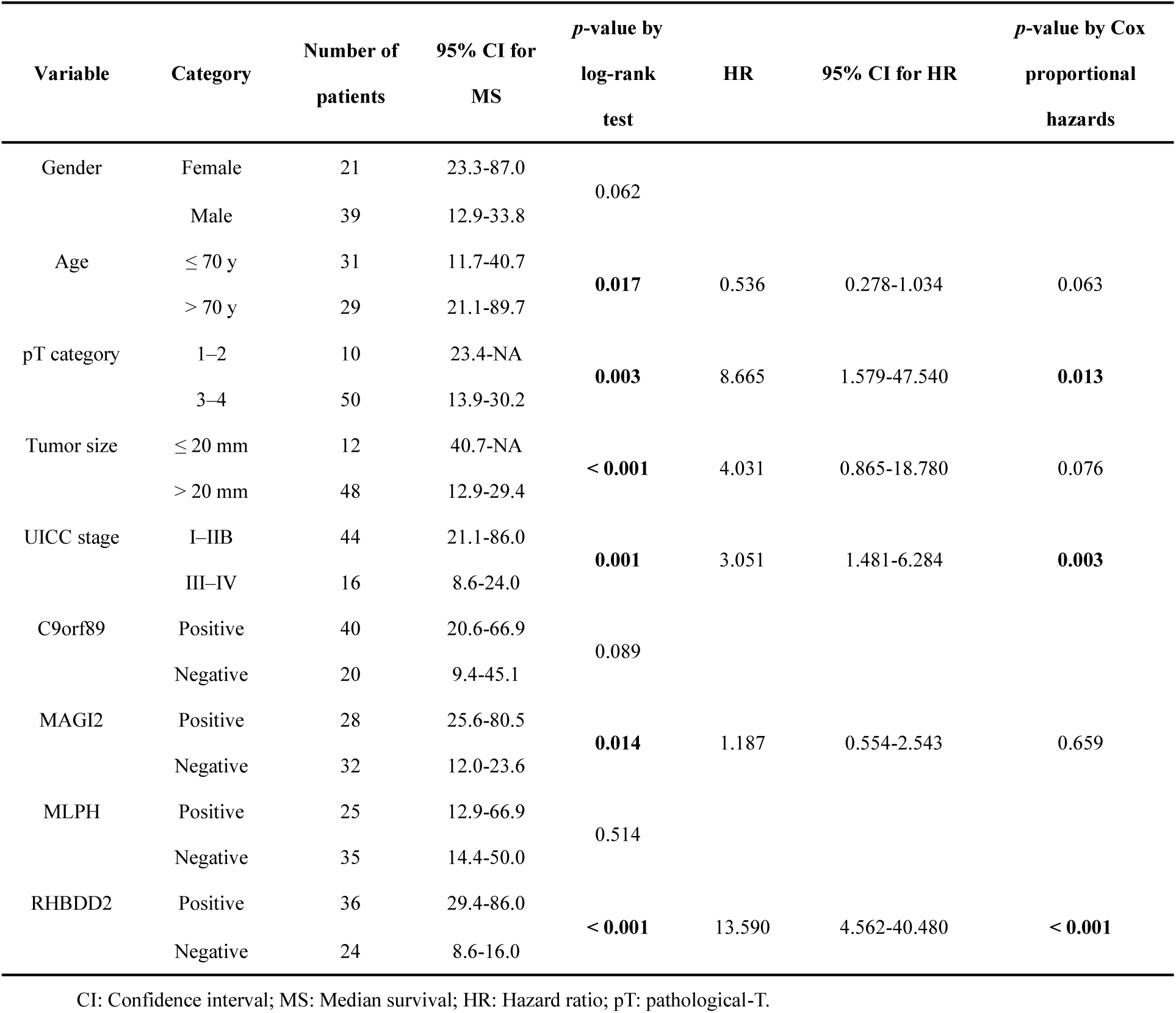
Univariate and multivariate analyses of factors associated with overall survival in patients with pancreatic ductal carcinoma

Additionally, immunohistochemical staining of CXCL12 was conducted to assess the association between RHBDD2 and CXCL12 expression in fibroblasts surrounding carcinoma cells. As a result, an inverse correlation was found between RHBDD2 and CXCL12 expression (Figure 5C).

## Discussion

In this study, Dendra2, which can convert its fluorescence wavelength by UV laser illumination, was introduced into adherent supporting cells combined with the CRISPR library and succeeded in isolating drug resistance inducible supporting cells close to tumor cells surviving and proliferating under anticancer drug exposure. This gene screening system, which focuses on labeling and isolating peritumoral responsible mutant cells, not selectively growing tumor cells, under microscopic observation, has not been reported previously and is a useful technique in comprehensive screenings of cell–cell interactions. However, it would be more efficient to search for candidate genes if single-cell RNA-seq was conducted for photoconverted cell clones without the expansion of targeted cells and the automation of laser illumination on the targeted cells under the fluorescence microscope.

*RHBDD2* was identified as a new gene responsible for drug resistance with cell–cell interactions, and clinical data also supported that the loss of RHBDD2 in supporting cells was suggested as a poor prognosis factor. *RHBDD2* is a member of the rhomboid family of membrane-bound proteases and is known to be overexpressed in the advanced stages of breast cancers and colorectal cancers (Abba et al., 2009) (Lacunza et al., 2012) (Palma et al., 2020). RHBDD2 is also reported that its overexpression promotes chemoresistance and invasive phenotypes of rectal cancer (Palma et al., 2020). However, these previous studies focused on the expression of RHBDD2 in the tumor cells themselves, and there are no reports examining its expression in peritumoral supporting cells. The pathway linking RHBDD2 and CXCL12 is a new finding in the present study, and the axis of RHBDD2–CXCL12 in cell–cell interactions could be a new drug target.

The drug resistance induced by the TME includes the prevention of drug absorption and the immune clearance of tumor cells (Vasan et al., 2019). Cell–cell adhesion systems in stromal cells, ECM, and cancer cells, such as integrin αvβ1 interactions, are known to convey such signals (Housman et al., 2014). Activating the tumor via secretory systems, such as the paracrine, juxtacrine, and autocrine, are also a mechanism by which cell–cell interactions in the microenvironment induce anticancer drug resistance, and CAF-derived secretion, such as through cytokines, chemokines, and growth factors, are known to be involved in tumor growth, progression, and drug resistance (Chen & Song, 2019) (Kanzaki & Pietras, 2020) (Sahai et al., 2020) (Wu et al., 2021) (Su et al., 2018). It is also known that PI3K-AKT-mTOR signaling pathway activation via CAF-derived secretion, such as through CXCL12, VCAM1, IL-22, CXCL5, HGF, and SPARC, induces tumor proliferation, migration, and stemness (Wu et al., 2021). In pancreatic cancer, elevated CXCL12 in CAFs is known to promote tumor progression via CXCL12–CXCR4 interaction (Li et al., 2016). In the present screening experiment, the cytokine signaling pathway in GSEA was up-regulated. *RHBDD2*-CKO-induced up-regulation of CXCL12 in stromal cells leads to anticancer drug resistance via activating the PI3K-Akt-mTOR pathway in tumor cells.

In recent years, many genetic screening methods that combine perturbation screening based on the CRISPR libraries with single-cell analysis, such as Perturb-seq, CITE-seq, and Perturb-CITE-seq, have been established, and they have led to the development of studies into the relationship between mutation and function on a single-cell basis (Dixit et al., 2016) (Datlinger et al., 2017) (Frangieh et al., 2021) (Datlinger et al., 2021) (Stoeckius et al., 2017). The characteristics of tumor cells with random mutations and their surrounding peritumoral stromal cells and inflammatory cells have been elucidated by applying spatial gene expression analysis technology, Visium, with CRISPR screening; however, this system screened the random mutation in the tumor cells and analyzed the expression profiles of stromal cells, i.e. not screened with random mutation in the stromal cells (Dhainaut et al., 2022). These techniques can analyze single supporting cells with randomly mutated tumors with CRISPR screening; however, it is difficult to isolate living supporting cells based on the distance and position of the tumors.

A novel screening method that combines the CRISPR library and photoconversion into cell–cell interactions was established. In future applications, CAFs/tumor-infiltrating lymphocytes (TILs) in actual tissues would be promising targets. This method could be useful to reveal new mechanisms behind all kinds of cell–cell interactions and collecting living cells is also a great advantage of biological analysis. It could also be a new platform for discovering the new targets of drugs in combination with conventional chemotherapy.

## Materials and Methods

### Key Resources Table

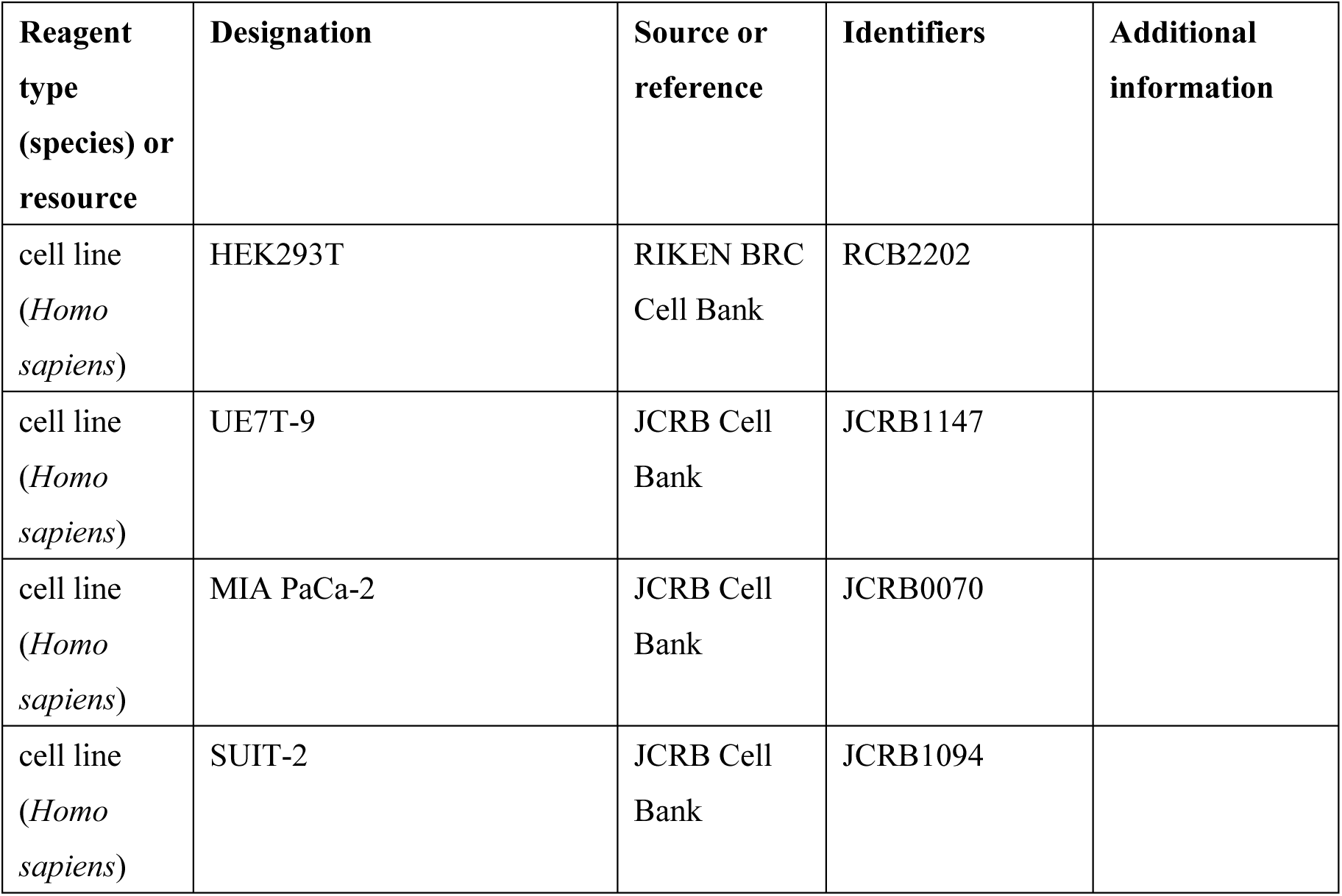
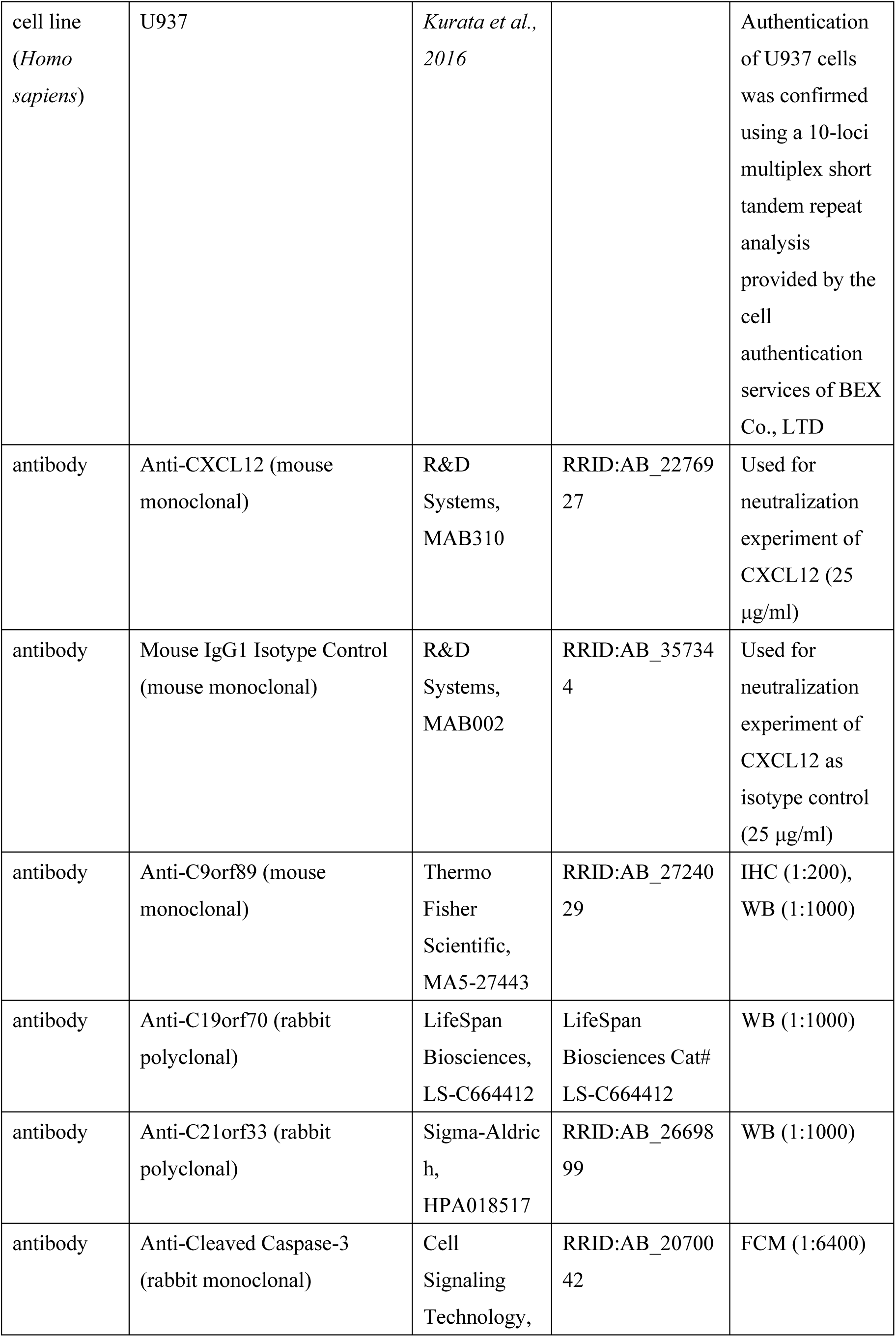
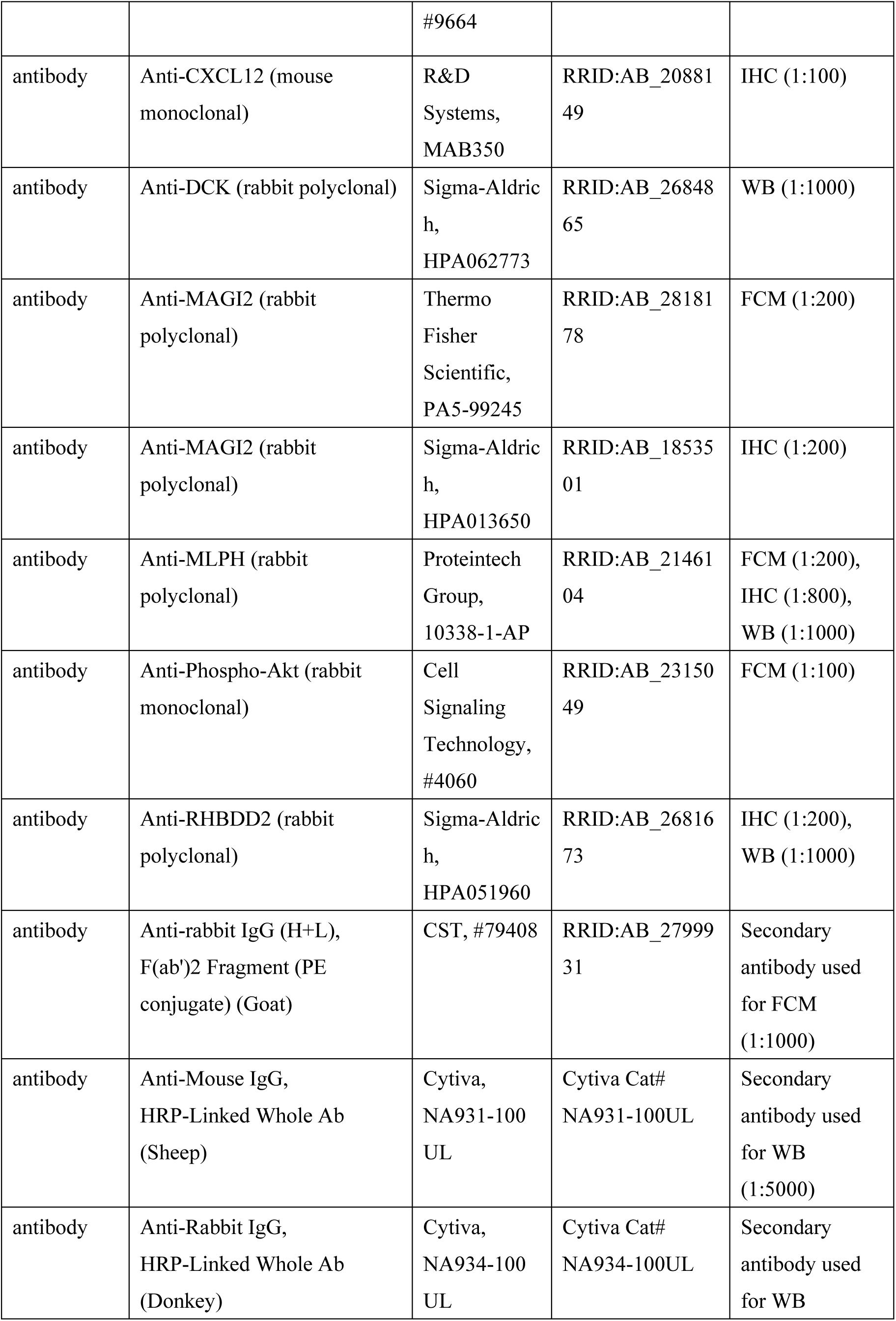
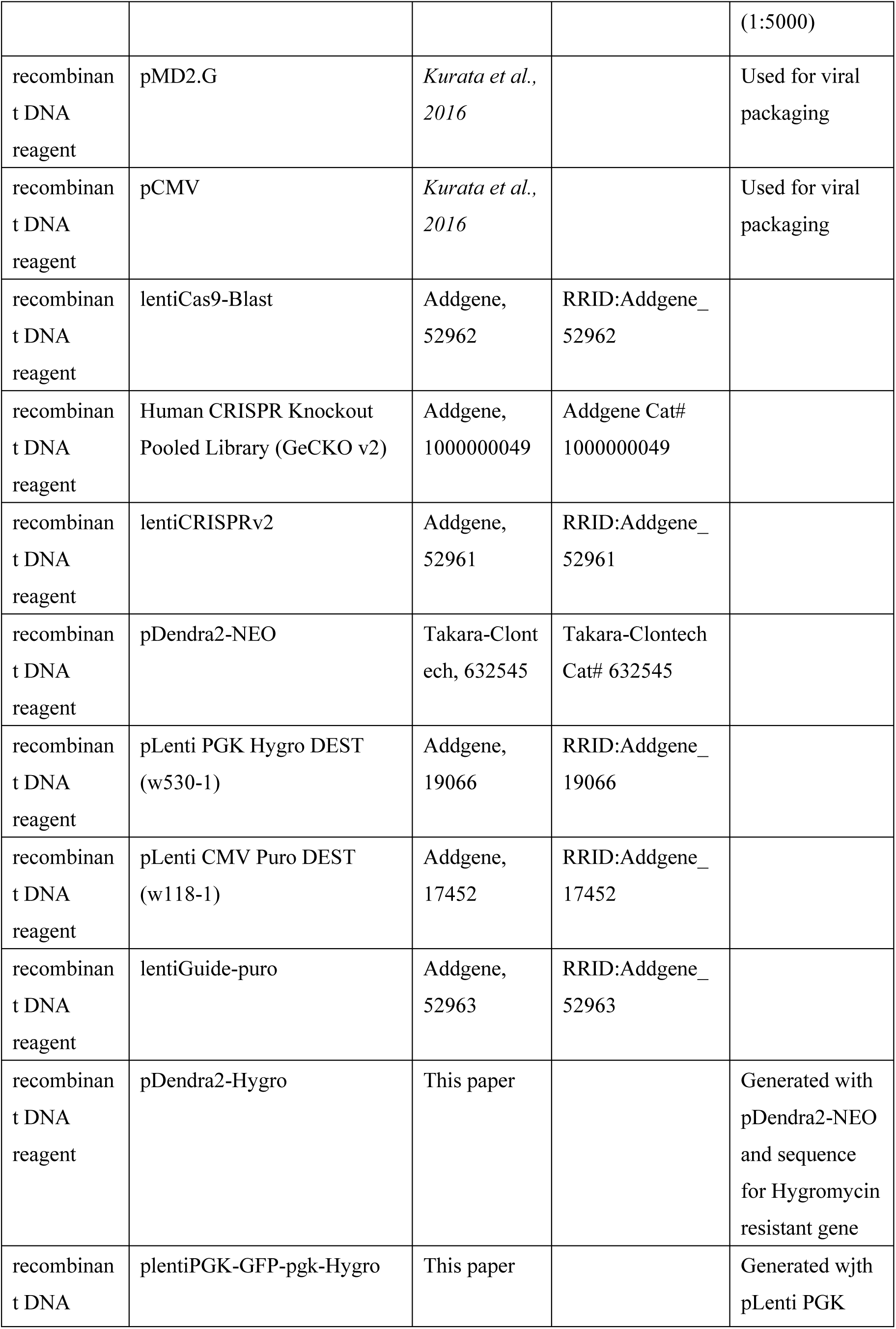
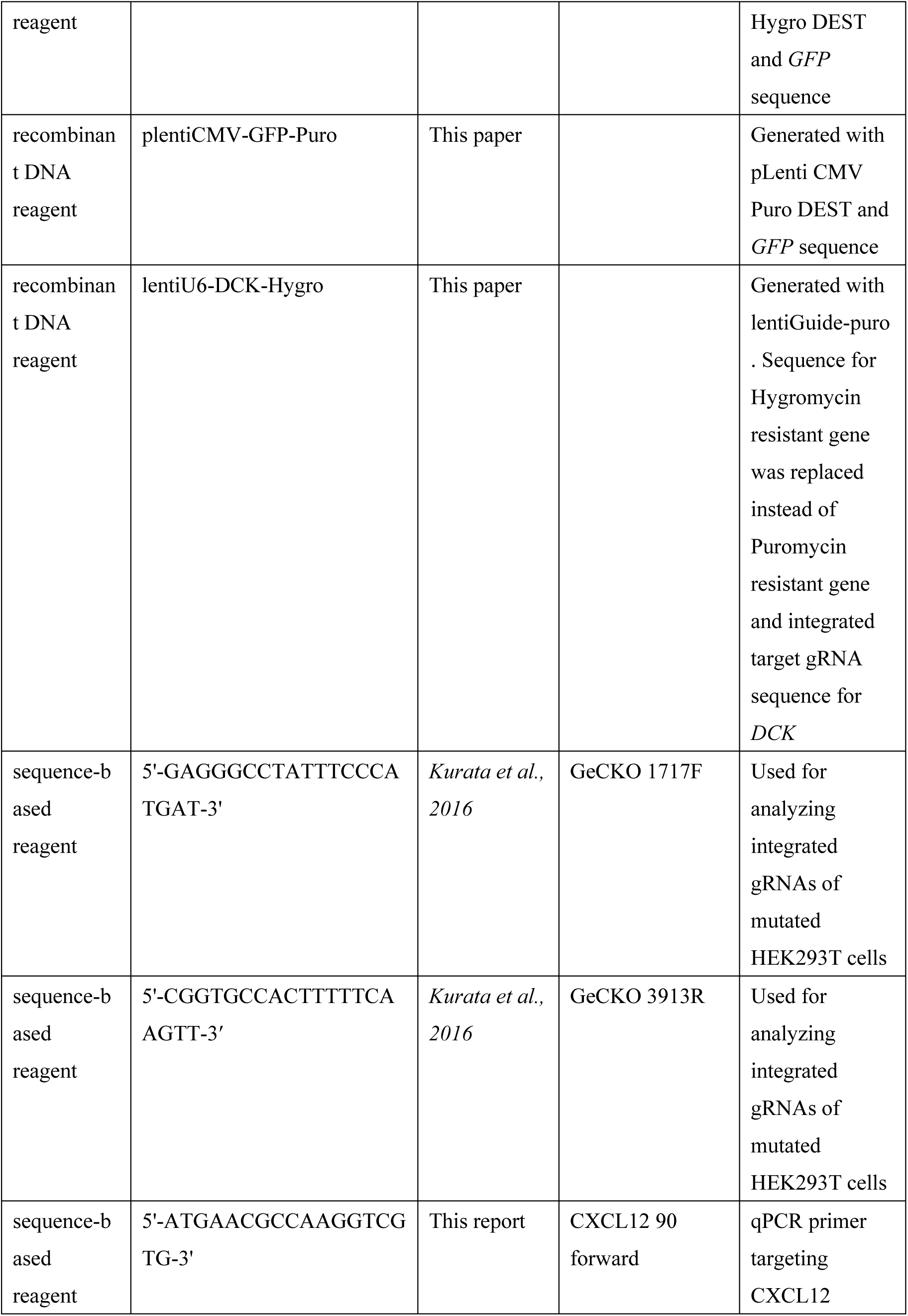
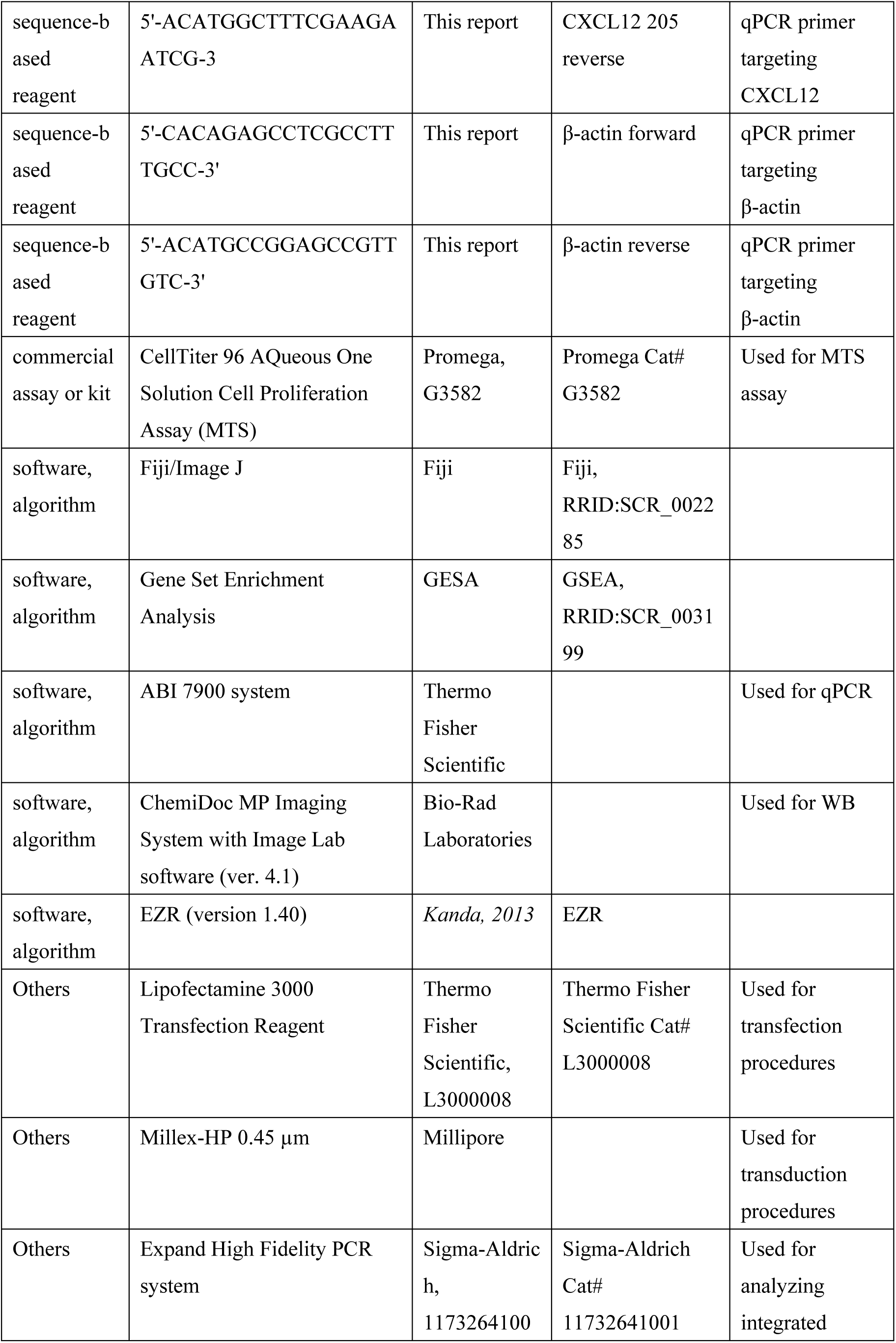
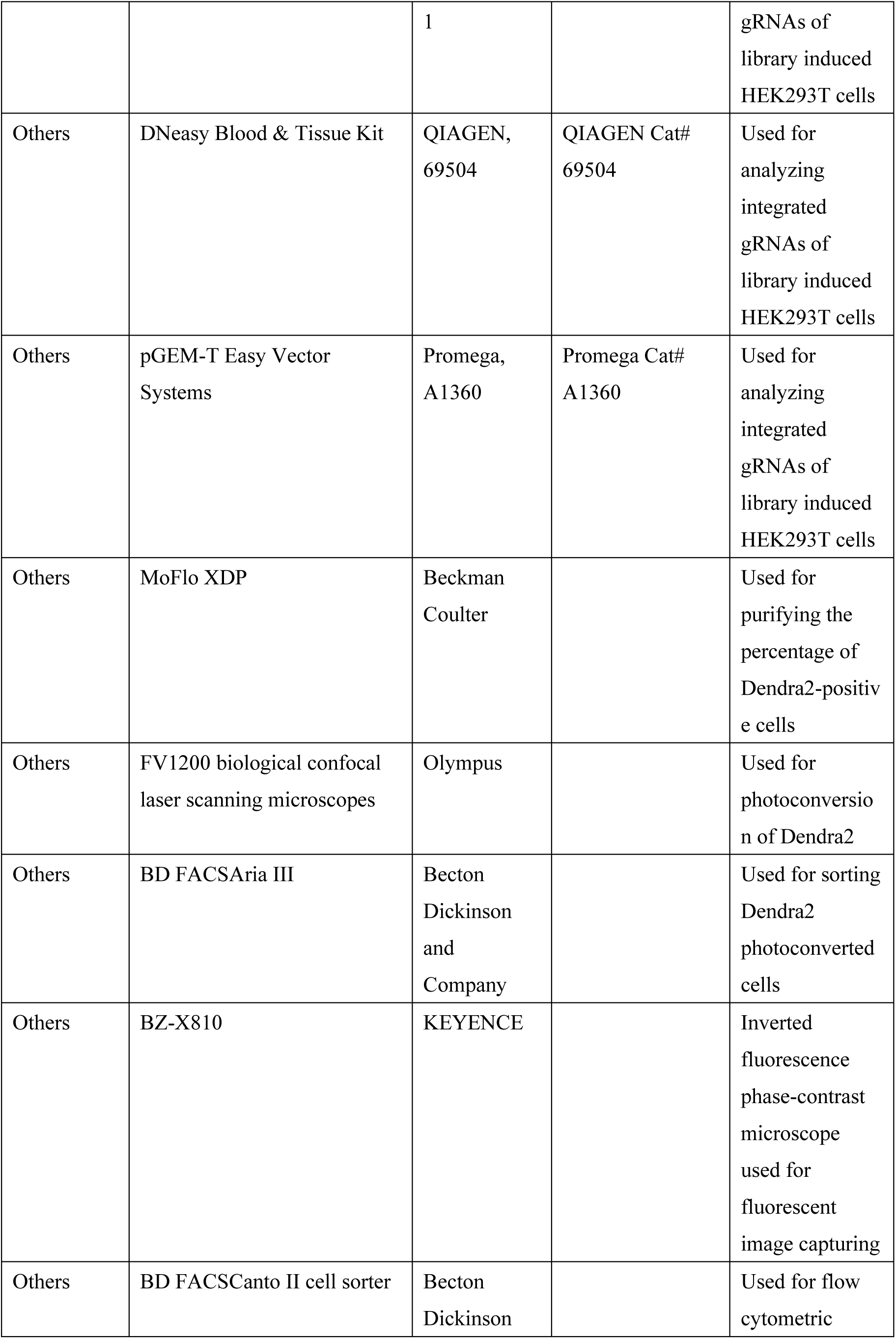
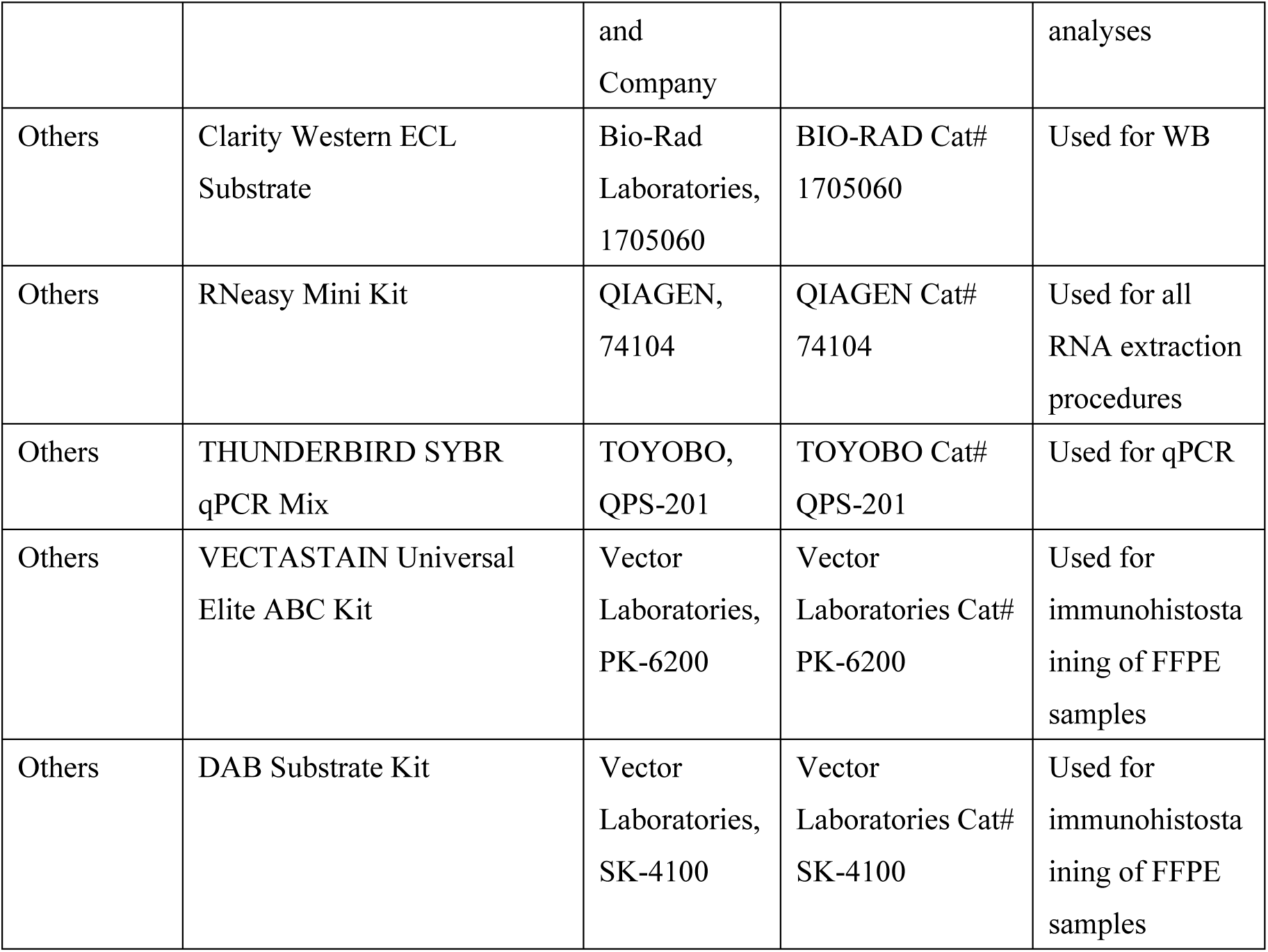

### Cell culture and transduction

All cell lines used in all experiments are listed in the Key resources table. HEK293T and UE7T-9 cells were maintained in Dulbecco’s modified Eagle’s medium (DMEM), U937 and SUIT-2 cells were maintained in Roswell Park Memorial Institute 1640 (RPMI), and MIA PaCa-2 cells were maintained in Eagle’s minimum essential medium (EMEM), respectively, with 10% fetal bovine serum (FBS) and 1% penicillin/streptomycin with 5% CO2 at 37 °C. Regarding the transduction of the lentiviruses, 5 × 10^5^ cells/well of HEK293T cells were seeded in a 6-well plate one day before the transfection. The next day, 3 µg of lentivirus plasmid was transfected with 1 µg of pMD2.G and 2 µg of pCMV using the Lipofectamine 3000 reagent. Twelve hours after transfection, the medium was changed to fresh DMEM. The virus supernatant was harvested at 48 h after transfection and then filtered with Millex-HP 0.45 µm. Then, 1 × 10^6^ cells/well of viral inducible cells were seeded in a 6-well plate one day prior to transduction and were transduced with this lentiviral supernatant with 5 µg/ml polybrene. Transduction was performed using spin infection followed by 30 min of centrifugation at 1,800 rpm and additional incubation for 2 h. Then, the supernatant was changed for the fresh culture medium. After a two-week selection with the antibiotics corresponding to resistance for each plasmid, cells were conducted in the following experiments.

HEK293T clones of *C9orf89*-CKO*, C19orf70*-CKO*, C21orf33*-CKO*, MAGI2*-CKO*, MLPH*-CKO, *RHBDD2*-CKO, and Non-target and UE7T-9 clones of *C9orf89*-CKO*, MAGI2*-CKO*, MLPH*-CKO, *RHBDD2*-CKO, and Non-target were established with the limiting dilution methods.

All plasmids used in all experiments are listed in the Key resources table.

### Indirect CRISPR screening

A vector pDendra2-Hygro was generated by integrating the hygromycin-resistant gene and P2A sequence into the multi-cloning site at the N-terminus of the pDendra2-N Vector. The lentiCas9-Blast was transduced to HEK293T cells and treated with blasticidin (10 µg/ml) for 2 weeks. The pDendra2-Hygro was induced by lipofection in Cas9-expressing HEK293T cells and treated with hygromycin B (200 µg/ml) for 2 weeks, and green fluorescence-positive cells were sorted using flow cytometry (MoFlo XDP) to purify the percentage of Dendra2-positive cells. The Human CRISPR Knockout Pooled Library A in lentiGuide-Puro was transduced into Cas9-Dendra2-expressing HEK293T cells and then treated with puromycin (1 µg/ml) for 2 weeks. The multiplicity of infection (MOI) was calculated and provided at 0.5. Library transduction was performed in two batches of 7.2 × 10^7^ cells in each trial and a total of 1.44 × 10^8^ cells of HEK293T. These 5 × 10^3^ cells/well of Dendra2 library-inducted HEK293T cells were conducted for screening in 15 96-well plates (for a total of 1,440 wells). Incubation HEK293T for 48 h to adhere to the bottoms of the plates, then 5 × 10^3^ cells of U937 were seeded per well, and an additional 48 h later, both U937 and the Dendra2 library-inducted HEK293T cells were exposed to 3 µM of cytarabine for 120 h. Regarding the wells where surviving U937 colonies were observed, both surviving U937 and Dendra2 library-inducted HEK293T cells were transferred to 24-well plates to expand the scale of Dendra2 library-inducted HEK293T, and co-culture was restarted with fresh U937 with cytarabine and then re-scaled up to 6-well plates. All supporting cells close to the viable U937 colonies under cytarabine exposure observed in 6-well plates were applied to photoconversion. Photoconversions were performed using FV1200 biological confocal laser scanning microscopes. The supporting cells close to the viable U937 colonies were identified by differential interference images generated by the FV1200. Observations and image captures of the green form of Dendra2 were generated using the 473 nm laser, whereas the red form was generated with the 559 nm laser. For photoconversion, the 405 nm laser at maximum power (100%) was used. The duration of the illumination period was set to 60 s, enough to observe the red form of Dendra2 under the 559 nm laser. The dotted lines showed the target area to be illuminated on the operating screen (Figure 1B, Step 4). Illumination was performed using the “bleach” mode of the FV1200 (Taniguchi et al., 2017). Dendra2 photoconverted HEK293T cells were sorted using the BD FACSAria III in a sterile environment. The cells were initially gated based on FSC-A and SSC-A channels to exclude the debris and dead cells.

Subsequently, the PI channel-positive cells were sorted as the populations that exhibited negligible signals in the un-photoconverted negative controls. Approximately 1,000 photoconverted cells were sorted per each well of the cells in the wells that underwent photoconversion.

The integrated gRNAs were sequenced by the following primers using the Expand High Fidelity PCR system (Sigma-Aldrich): GeCKO 1717F 5′-GAGGGCCTATTTCCCATGAT-3′ and GeCKO 3913R 5′- CGGTGCCACTTTTTCAAGTT -3′. Genomic DNA isolations from each well were acquired using a DNeasy Blood & Tissue Kit. The fragments were sequenced by TA cloning with pGEM-T Vector Systems. The analyzed sequences of the gRNA region were compared with a list of human library A gRNA sequences (https://media.addgene.org/cms/filer_public/a4/b8/a4b8d181-c489-4dd7-823a-fe267fd7b277/human_geckov2_library_a_09mar2015.csv) to identify candidate genes.

### Cell co-culture validation experiments

The sequences of all oligonucleotides used in the experiments are provided in Table 1 and Supplementary File 8. The designed oligonucleotides were purchased from Invitrogen. Each oligonucleotide of all candidate genes and non-targeted gRNA (Non-target) were integrated into lentiCRISPRv2 plasmids following the manufacturer’s protocol.

HEK293T or UE7T-9 were transduced with lentiCRISPRv2 of each candidate gene. In subsequent experiments, HEK293T or UE7T-9 cells transduced with lentiCRISPRv2 Non-target were used as controls. After a two-week selection with puromycin, cells were conducted in the cell co-culture validation experiments. For labeling for the indication of live tumor cells, plentiPGK-GFP-pgk-Hygro was inducted to U937, and plentiCMV-GFP-Puro was inducted to MIA PaCa-2 and SUIT-2. LentiU6-DCK-Hygro was generated with lentiGuide-puro by replacing the sequence of the puromycin-resistant gene with the hygromycin-resistant gene and the integrated target gRNA sequence for DCK (Supplementary File 8 and previous report (Kurata et al., 2016)).

LentiU6-DCK-Hygro was transduced to Cas9-HEK293T. The HEK293T *DCK*-CKO clone was established with limiting dilution methods and then transduced with lentiCRISPRv2-gRNA to make knockout HEK293T cells of each candidate gene.

In the co-culture experiments of mutant-HEK293T with U937 or mutant-UE7T-9 with U937 cells, 1 × 10^5^ cells of the target mutated HEK293T or target mutated UE7T-9 cells were seeded in a 24-well plate with the RPMI medium. After 48 h of incubation, 1 × 10^5^ cells of GFP-positive U937 were added per well. After an additional 48 h of co-culture to make sure all cells could contact each other, cells were exposed to 5 μM of cytarabine for 48 h.

In the co-culture experiments of mutant-HEK293T with pancreatic cancer cells, 1 × 10^5^ cells of *DCK* and target mutated HEK293T cells and 1 × 10^5^ cells of GFP-positive pancreatic cancer cells (MIA PaCa-2 or SUIT2) were seeded in 24-well plates and co-cultured (with the EMEM medium for MIA PaCa-2 or the RPMI medium for SUIT-2). After 48 h of co-culture, cells were exposed to gemcitabine (MIA PaCa-2: 10 μM, SUIT-2: 3 μM) for 168 h.

For each co-culture experiment, three wells were prepared with the same conditions, and three fields were randomly captured for each well with the drug exposure. Fluorescent image capture was performed using an inverted fluorescence phase-contrast microscope BZ-X810. The field of view was 20× objective using a green laser. The count of viable cells per field of view was performed using Image J (https://imagej.net/software/fiji/) (Schindelin et al., 2012). To compare viability, the number of viable cells relative to that co-cultured with the control (Non-target) of supporting cells was calculated.

### The analysis of cleaved caspase-3 in the co-culture system

In the analysis of cleaved caspase-3 in the HEK293T-U937 co-culture system, 5 × 10^5^ HEK293T-CKO cells were seeded in 6-well plates. After 48 h of incubation, 5 × 10^5^ cells of U937 were added per well. After an additional 48 h of co-culture, cells were exposed to 5 μM of cytarabine for 48 h. As a control, HEK293T mono-culture groups were also conducted in the experiment. After 48 h of cytarabine exposure, all the cells in each well were collected, fixed with 4% paraformaldehyde, and permeabilized with 90% methanol following the manufacturer’s protocol. All collected cells were incubated for 1 h with anti-cleaved caspase-3 primary antibodies at room temperature, washed twice with PBS, and incubated with anti-rabbit IgG (H+L), F(ab’)2 Fragment (PE conjugate) for 30 min at room temperature and protected from light. Then, they were washed twice with PBS. The percentages of cleaved caspase-3-positive cells were evaluated using the BD FACSCanto II cell sorter. The cells were gated based on PE-A and SSC-A channels as populations that exhibited negligible signals in the unstained negative controls.

Information about all antibodies used in all experiments is provided in the Key resources table.

### The analysis of cleaved caspase-3 with the neutralization of CXCL12

In the experiments on the neutralization of CXCL12, 1 × 10^5^ cells of HEK293T *RHBDD2*-CKO or Non-target were seeded in 350 μl of fresh RPMI medium in a 24-well plate. After 48 h of incubation, 1 × 10^5^ cells of U937 were added in 350 μl of fresh RPMI medium with human/mouse CXCL12/SDF-1 antibody or mouse IgG1 isotype control (25 μg/ml). After 48 h of co-culture, cells were exposed to 5 μM of cytarabine for 48 h. The process used after the cytarabine exposure was the same as described in the cleaved caspase-3 detection in the co-culture system.

### The analysis of phospho-Akt

In the analysis of phospho-Akt of U937 co-cultured with HEK293T, U937 was labeled with GFP to evaluate phospho-Akt in U937 cells only. Then, 5 × 10^5^ cells of HEK293T *RHBDD2*-CKO or Non-target cells were seeded in 2 ml of RPMI in a 6-well plate. After 48 h of incubation, all mediums were changed with 5 × 10^5^ cells of GFP-positive U937 in 2 ml of RPMI medium. As a control, U937 mono-culture groups were also conducted in the experiment. After 48 h of co-culture, all cells in each well were collected, fixed with 4% paraformaldehyde, and permeabilized with 0.1% of Triton X-100 following the manufacturer’s protocol. All collected cells were incubated for 1 h with anti-phospho-Akt primary antibodies at room temperature, and the process used after the primary antibodies was the same as described in the cleaved caspase-3 detection. The expression of phospho-Akt of U937 was evaluated using the BD FACSCanto II cell sorter. The cells were gated based on FSC-A and SSC-A channels to exclude debris and dead cells, and they were further gated based on the GFP-A channel to exclude HEK293T. Subsequently, the cells were gated based on the PE-A channel as the populations that exhibited negligible signals in the unstained negative controls.

### RNA sequencing and analysis

Total RNA was extracted from HEK293T cells of *C9orf89*-CKO, *MAGI2*-CKO, *MLPH*-CKO, *RHBDD2*-CKO, and Non-target incubated in three wells of 6-well plates using the RNeasy Mini Kit following the manufacturer’s protocol, including the DNase I treatment. All RNAs were analyzed by the RNA sequencing service provided by Novogene Co., Ltd. (Beijing, China) using the NovaSeq 6000 Sequencing System (Illumina, San Diego, CA, USA) and 6 Gb of the Eukaryotic mRNA Library. The RNA profile data were deposited in a public database, Gene Expression Omnibus (GEO), under access number GSE 203256. For GSEA, normalized expression data were analyzed and visualized with GSEA software (version 4.2.0, https://www.gsea-msigdb.org/gsea/index.jsp).

The normalized enrichment score (NES), nominal *p*-value (NOM-*p*), and false discovery rate *q*-value (FDR-*q*) were calculated for comparison, and the categories selected were universally up-regulated in HEK293T cells of *C9orf89*-CKO, *MAGI2*-CKO, *MLPH*-CKO, and *RHBDD2*-CKO. The relative enrichment of individual genes was assessed based on the rank metric score following the GSEA. Volcano plots were generated using Novosmart software (Novogene).

A quantitative polymerase chain reaction (qPCR) was performed based on SYBR Green, and the expression of *CXCL12* was normalized to β-actin levels. Primers for *CXCL12* or β-actin are shown in the Key resources table. All assays were performed on an ABI 7900 system and analyzed with ABI 7900 SDS software v.2.4.1.

### Immunohistostaining of clinical samples

Sixty clinical samples of invasive pancreatic ductal carcinoma were obtained from 60 patients who underwent operations without neoadjuvant chemotherapy at the Tokyo Medical and Dental University Hospital, Tokyo, between 2008 and 2016. Specimens were obtained by surgical resection, fixed in 10% neutralized formalin, and embedded in paraffin according to routine protocols used in conventional histopathological examinations. Informed consent was obtained from all patients, and the study was approved by the ethics committees of Tokyo Medical and Dental University. All procedures were performed following the ethical standards established by these committees (M2000-1458-07).

Formalin-fixed, paraffin-embedded (FFPE) tissues were sectioned at a thickness of 4 μm, placed on silane-coated slides, and deparaffinized. Heat-based antigen retrieval, endogenous peroxidase blockade using 3% hydrogen peroxide, and blocking were performed with normal sera. The sections were incubated overnight with primary antibodies against C9orf89, CXCL12, MAGI2, MLPH, and RHBDD2 at 4 °C. Primary antibodies were detected using an ABC kit. Color development was performed using diaminobenzidine (DAB). The scoring of immunostainability was confirmed by two pathologists (K. Sugita and M. Kurata) based on the following criteria: not stained in fibroblasts was (-), positive in only a few fibroblasts or weakly positive were (1+), and strongly positive on most of the fibroblasts was (2+). Representative images of each antibody are shown. Based on scoring, (-) and (1+) were classified as negative groups, and (2+) was the positive group (Figure 5—figure supplement 1).

The following five parameters were evaluated as clinicopathologic factors: sex (female vs. male), age (≤ 70 y vs. > 70 y), pT category (1–2 vs. 3–4), tumor size (≤ 20 mm vs. > 20 mm), and UICC stage (I–IIB vs. III–IV).

### Cell viability assessment

Cell viability under drug exposure was assessed using the MTS assay following the manufacturer’s protocol.

In the analysis of cell sensitivities to cytarabine or gemcitabine, 1 × 10^3^ cells were seeded in a96-well plate for 24 h and then treated with cytarabine or gemcitabine with serial dilution methods. After 48 h of drug exposure, cell viability was evaluated with an MTS assay, and IC50 was estimated.

In the analysis of U937 cell viability cultured with the supernatant from HEK293T, 1 × 10^6^ of HEK293T *RHBDD2*-CKO were seeded in 2 ml of RPMI in a 6-well plate. After 48 h of cells incubation, the cultured supernatant was collected and filtered. Then, 3 × 10^4^ cells of U937 were seeded in 150 μl of “collected supernatant of HEK293T *RHBDD2*-CKO” in a 96-well plate and incubated for 48 h, then treated with 250 nM of cytarabine. After 48 h of cytarabine exposure, cell viability was evaluated using the MTS assay.

### Confirmation of knockouts of candidate genes in CKO clones

To confirm the knockouts of candidate genes in HEK293T-CKO clones or UE7T-9 CKO clones, western blot analysis or flow cytometric analysis were performed. Western blot analysis was applied for HEK293T cells of *DCK*-CKO, *C9orf89*-CKO, *C19orf70*-CKO, *C21orf33*-CKO, and *RHBDD2*-CKO, as well as UE7T-9 cells of *C9orf89*-CKO, *MLPH*-CKO, and *RHBDD2*-CKO. In the western blot analysis, cells were lysed and sonicated in the SDS-PAGE sample buffer. Protein samples were separated by SDS-PAGE and then transferred to PVDF membranes. β-actin was used as the loading control. Anti-rabbit or mouse HRP-linked IgG antibody was used as a secondary antibody. Detection was performed using Clarity Western ECL Substrate. The blot was imaged using the ChemiDoc MP Imaging System with Image Lab software ver. 4.1.

Flow cytometric analysis was applied to HEK293T cells of *MAGI2*-CKO and *MLPH*-CKO, as well as the UE7T-9 cell of *MAGI2*-CKO owing to their difficulty of detection with western blot in the parental cells. The process used before the flow cytometric analysis was the same as described in the cleaved caspase-3 detection. In the flow cytometric analysis, cells were gated based on FSC-A and SSC-A to exclude debris and dead cells. Subsequently, cells were evaluated with PE signals by comparing them with stained control cells and unstained negative controls.

## Statistical analysis

One-way analysis of variance (ANOVA) with Dunnett’s test or Bonferroni’s test was used for the analysis of data involving multiple groups. The two-tailed unpaired Student’s t-test was used to compare the two groups. OS was determined from the date of diagnosis and that of last follow-up or death. Kaplan–Meier survival curves were used to estimate rates of OS. The log-rank test was used to analyze differences in survival between the groups. Univariate and multivariate analyses were performed using the Cox proportional hazard regression model. Correlations between the two groups were assessed using Fisher’s exact test. All *p-*values < 0.05 were considered statistically significant.

In GSEA, results were considered significant if both NOM-*p* < 0.05 and FDR-*q* < 0.25 were satisfied. In all graphs, the data are presented as mean ± standard deviation (SD). Meanwhile, the results were obtained independently in triplicate for each experiment, and the experiments were usually repeated two or three times. All statistical analyses were performed using the free statistical software EZR (version 1.40) (Kanda, 2013).

## Acknowledgements

The authors would like to thank Soichiro Yoshida from the Department of Urology, Graduate School of Medical and Dental Sciences, Tokyo Medical and Dental University, and the members of the stem cell laboratory of Tokyo Medical and Dental University for their technical assistance and advice.

## Author contributions

Keisuke Sugita, Methodology, Investigation, Validation, Formal analysis, Data curation, Visualization, Writing – original draft preparation; Iichiroh Onishi, Supervision, Writing – review & editing; Ran Nakayama, Investigation, Validation, Formal analysis, Data curation, Writing – review & editing; Sachiko Ishibashi, Investigation, Formal analysis, Data curation, Writing – review & editing; Masumi Ikeda, Investigation, Writing – review & editing; Miori Inoue, Investigation, Writing – review & editing; Rina Narita, Investigation, Validation, Formal analysis, Visualization, Writing – review & editing; Shiori Oshima, Investigation, Validation, Formal analysis, Writing – review & editing; Kaho Shimizu, Investigation, Validation, Formal analysis, Writing – review & editing; Shinichiro Saito, Investigation, Validation, Writing – review & editing; Shingo Sato, Methodology, Resources, Supervision, Writing – review & editing; Branden S. Moriarity, Investigation, Resources, Supervision, Writing – review & editing; Kouhei Yamamoto, Supervision, Writing – review & editing; David A. Largaespada, Resources, Supervision, Writing – review & editing; Masanobu Kitagawa, Supervision, Project administration, Writing – review & editing; Morito Kurata, Conceptualization, Methodology, Formal analysis, Investigation, Resources, Data curation, Project administration, Funding acquisition, Supervision, Writing – original draft preparation.

## Data Availability

Original data of western blots, raw data of flow cytometry, and plasmid maps in the present study have been deposited in figshare.com.

Figure 1B,C (10.6084/m9.figshare.19769062).

Figure 3—figure supplement 1 (10.6084/m9.figshare.19771483).

Figure 4A (10.6084/m9.figshare.19771147).

Figure 4E (10.6084/m9.figshare.19771156).

Figure 4F (10.6084/m9.figshare.19771555).

Figure 4—figure supplement 2 (10.6084/m9.figshare.19771504).

Plasmid maps (10.6084/m9.figshare.19771507).

RNA-seq data have been deposited in Gene Expression Omnibus (GEO) under access number GSE 203256.

---

**Figure 1—figure supplement 1.**
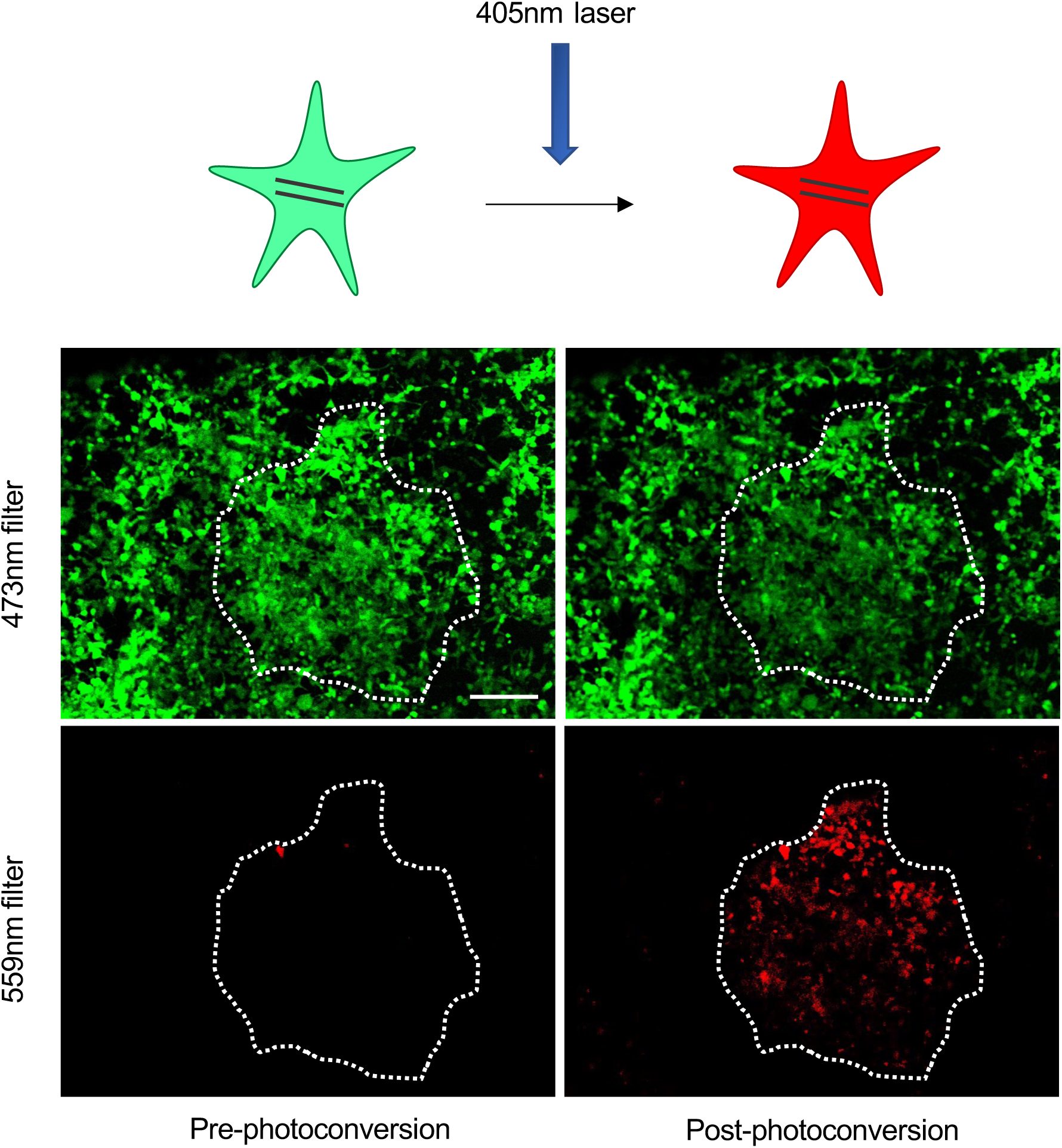
Dendra2 photoconversion. Dendra2 is a green fluorescent protein, and illumination with the 405 nm UV laser can irreversibly convert its fluorescence wavelength from green to red. The lower panel shows photographs of the actual cells pre- and post-photoconversion, in which red fluorescence was induced by illuminating the dotted line with the 405 nm laser. Image captures of the green form of Dendra2 were performed with the 473 nm laser (upper tier), whereas the red form was performed with the 559 nm laser (lower tier). Scale bar, 50 μm.

**Figure 2—figure supplement 1.**
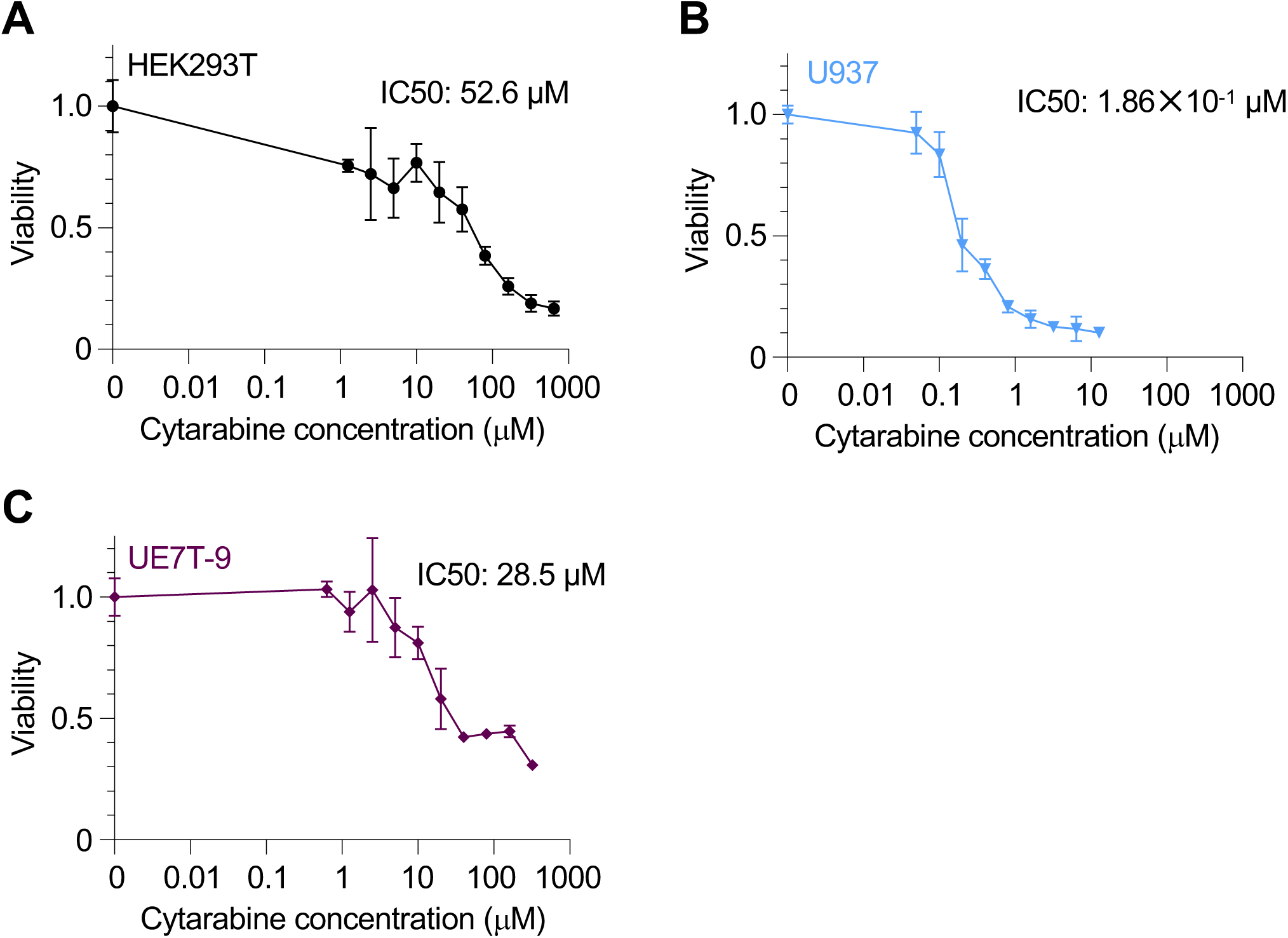
Cell viabilities under cytarabine exposure. (**A**-**C**) Dose-response curves of HEK293T (A), U937 (B), and UE7T-9 (C) cells treated with cytarabine for 48 h with biological triplication. IC50 to cytarabine are shown in each graph. Data are represented as mean ± SD. Figure 2**—figure supplement 1—source data 1.** Values for the graph in Figure 2—figure supplement 1A. Figure 2**—figure supplement 1—source data 2.** Values for the graph in Figure 2—figure supplement 1B. Figure 2**—figure supplement 1—source data 3.** Values for the graph in Figure 2—figure supplement 1C.

**Figure 2—figure supplement 2.**
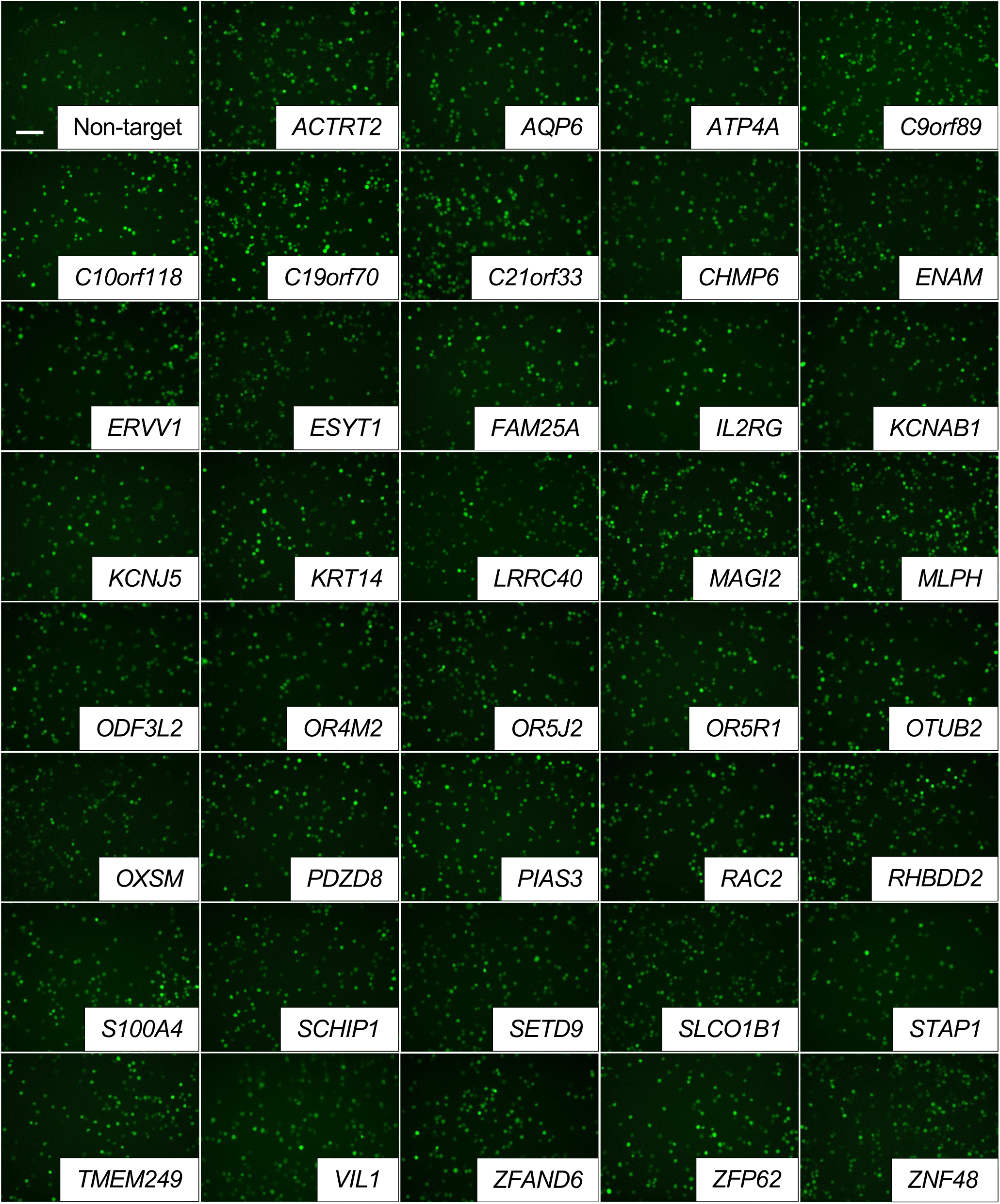
Representative images of GFP-positive cells in knockout mutant HEK293T-U937 co-culture experiments for each candidate. GFP-positive cells were indicated as viable U937 cells. After 48 h of exposure to cytarabine, images were captured under a green laser with a 20x objective field of view. Scale bar, 50 μm.

**Figure 2—figure supplement 3.**
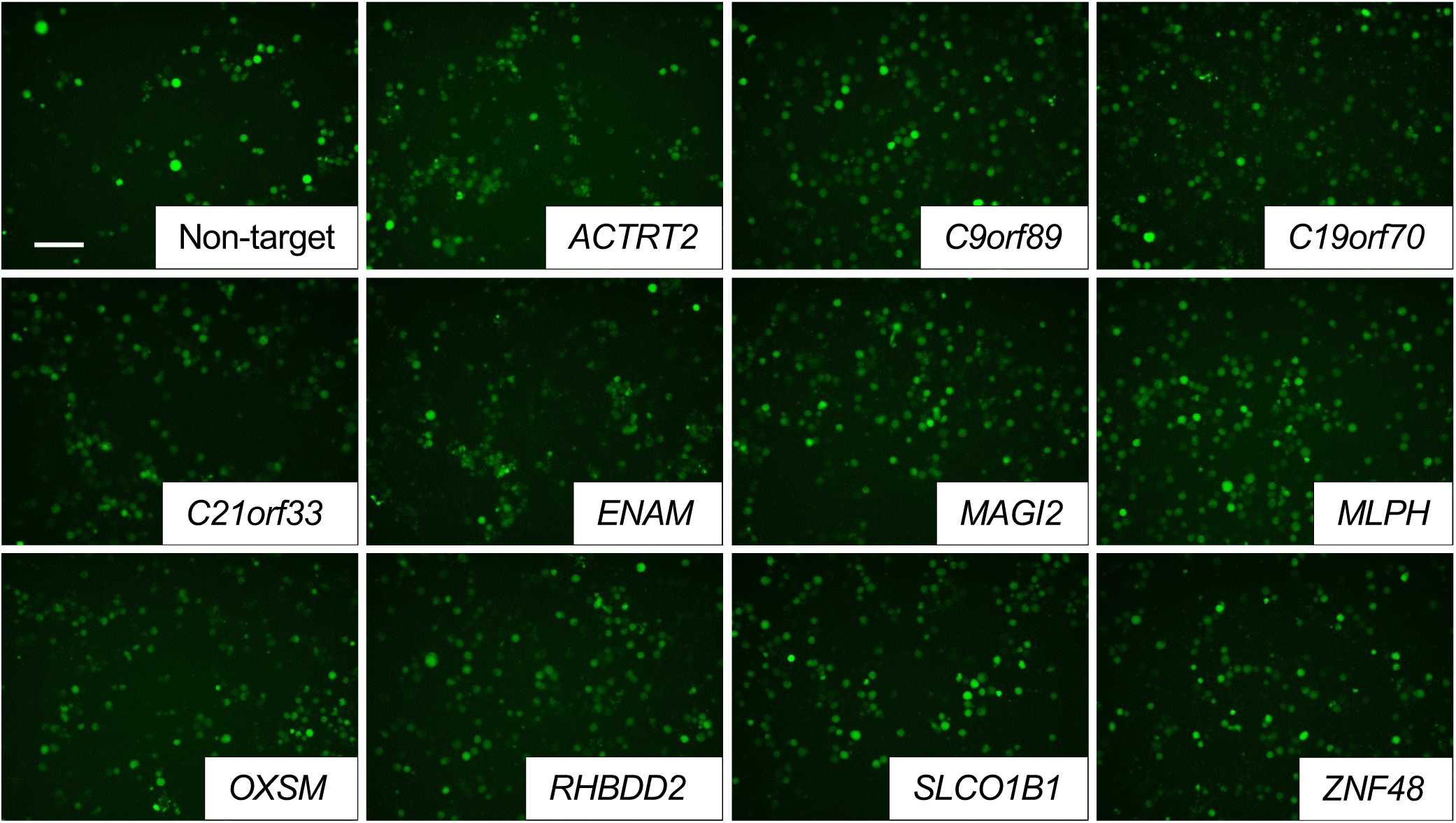
Representative images of GFP-positive cells in knockout mutant UE7T-9-U937 co-culture experiments for each candidate. GFP-positive cells were indicated as viable U937 cells. After 48 h of exposure to cytarabine, images were captured under a green laser with a 20x objective field of view. Scale bar, 50 μm.

**Figure 3—figure supplement 1.**
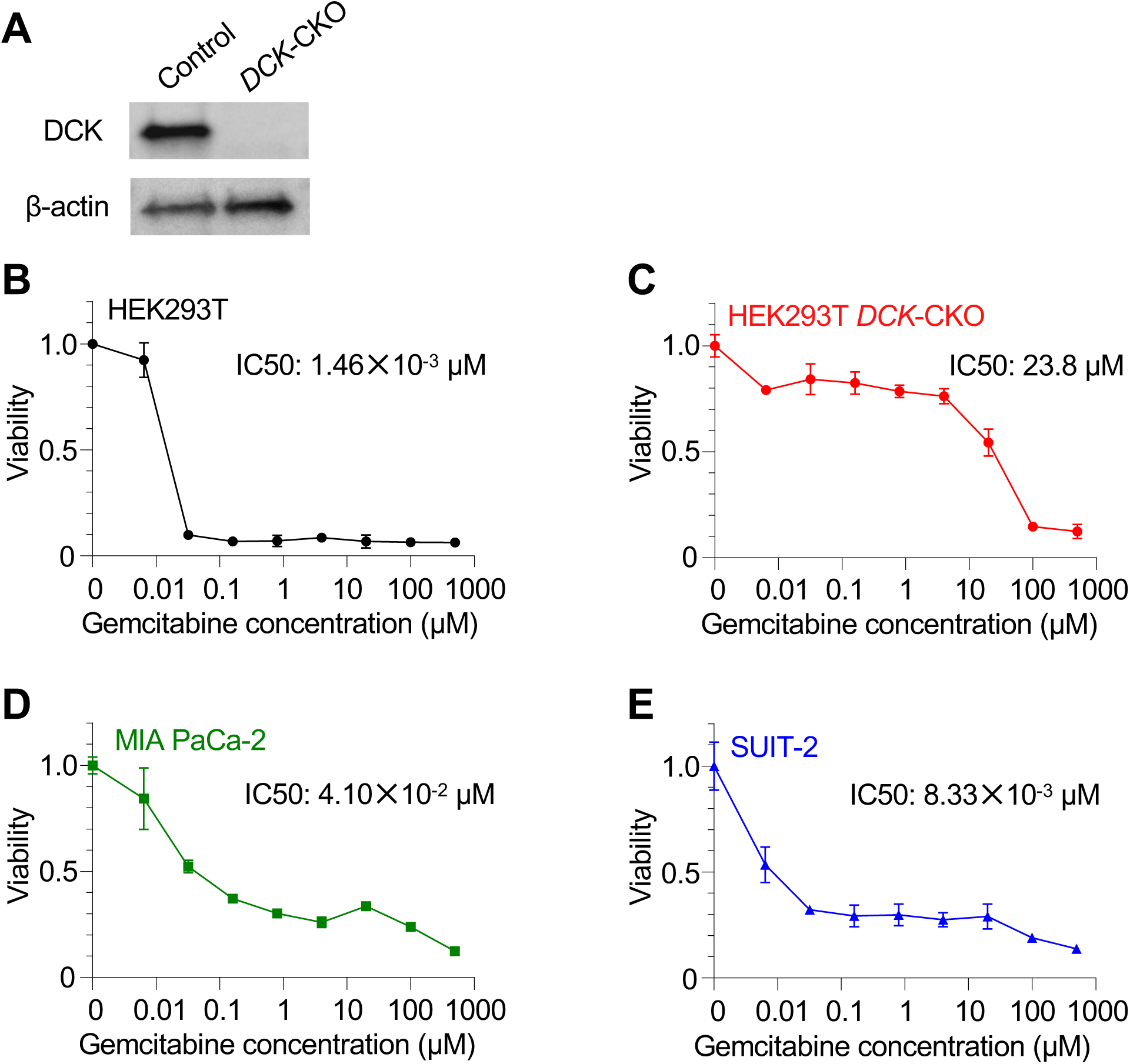
Validation experiments to determine drug resistance with cell– cell interactions in HEK293T-pancreatic cancer cell co-culture experiment. (**A**) Loss of DCK in HEK293T *DCK*-CKO clone in the western blot. (**B–E**) Dose-response curves of HEK293T (B), HEK293T *DCK-*CKO (C), MIA PaCa-2 (D), and SUIT-2 (E) cells treated with gemcitabine for 48 h with biological triplication. IC50 to gemcitabine is shown in each graph. Data are represented as mean ± SD. Figure 3**—figure supplement 1—source data 1.** Values for the graph in Figure 3—figure supplement 1B. Figure 3**—figure supplement 1—source data 2.** Values for the graph in Figure 3—figure supplement 1C. Figure 3**—figure supplement 1—source data 3.** Values for the graph in Figure 3—figure supplement 1D. Figure 3**—figure supplement 1—source data 4.** Values for the graph in Figure 3—figure supplement 1E.

**Figure 3—figure supplement 2.**
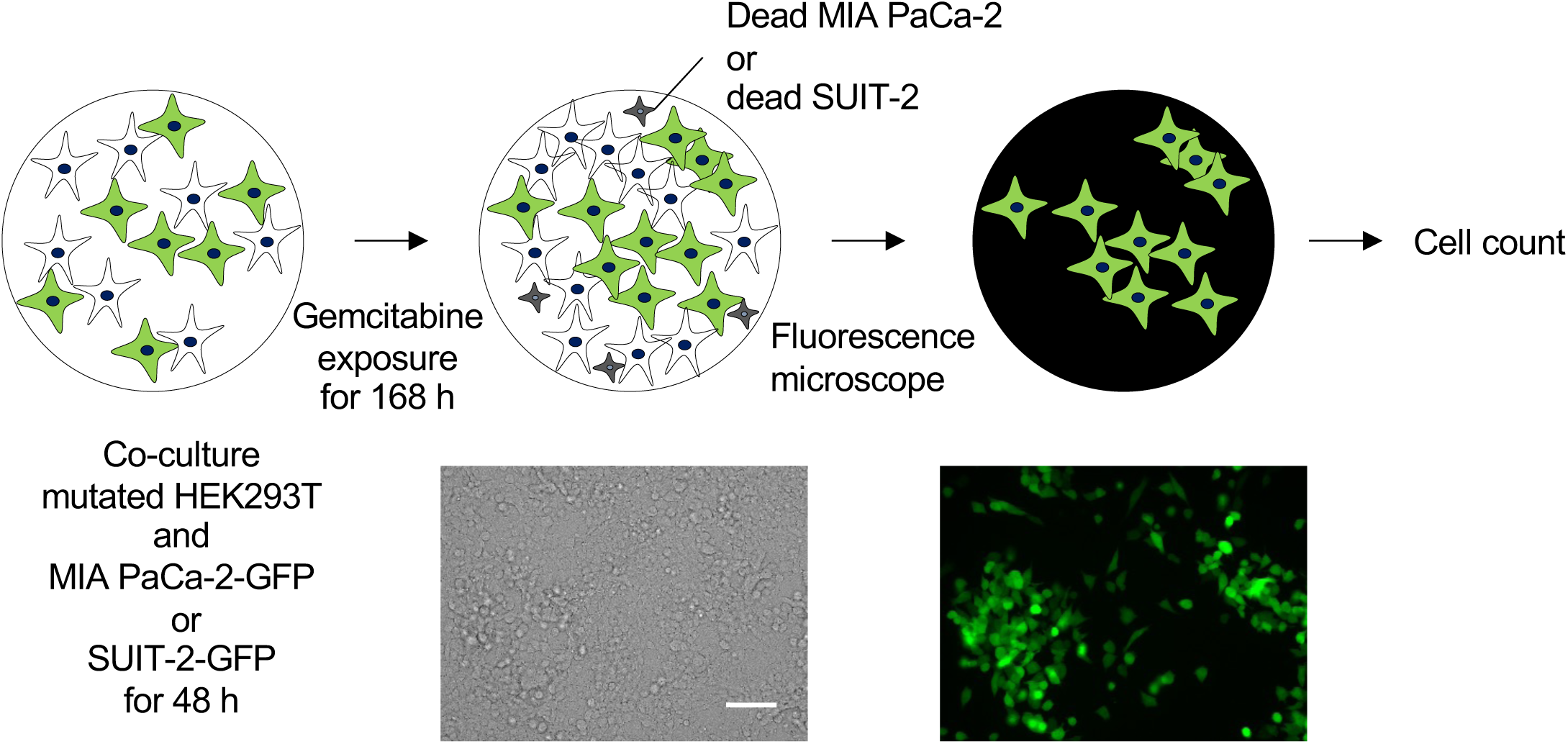
Experimental scheme of HEK293T-pancreatic cancer cell co-culture experiment. Scale bar, 50 μm.

**Figure 3—figure supplement 3.**
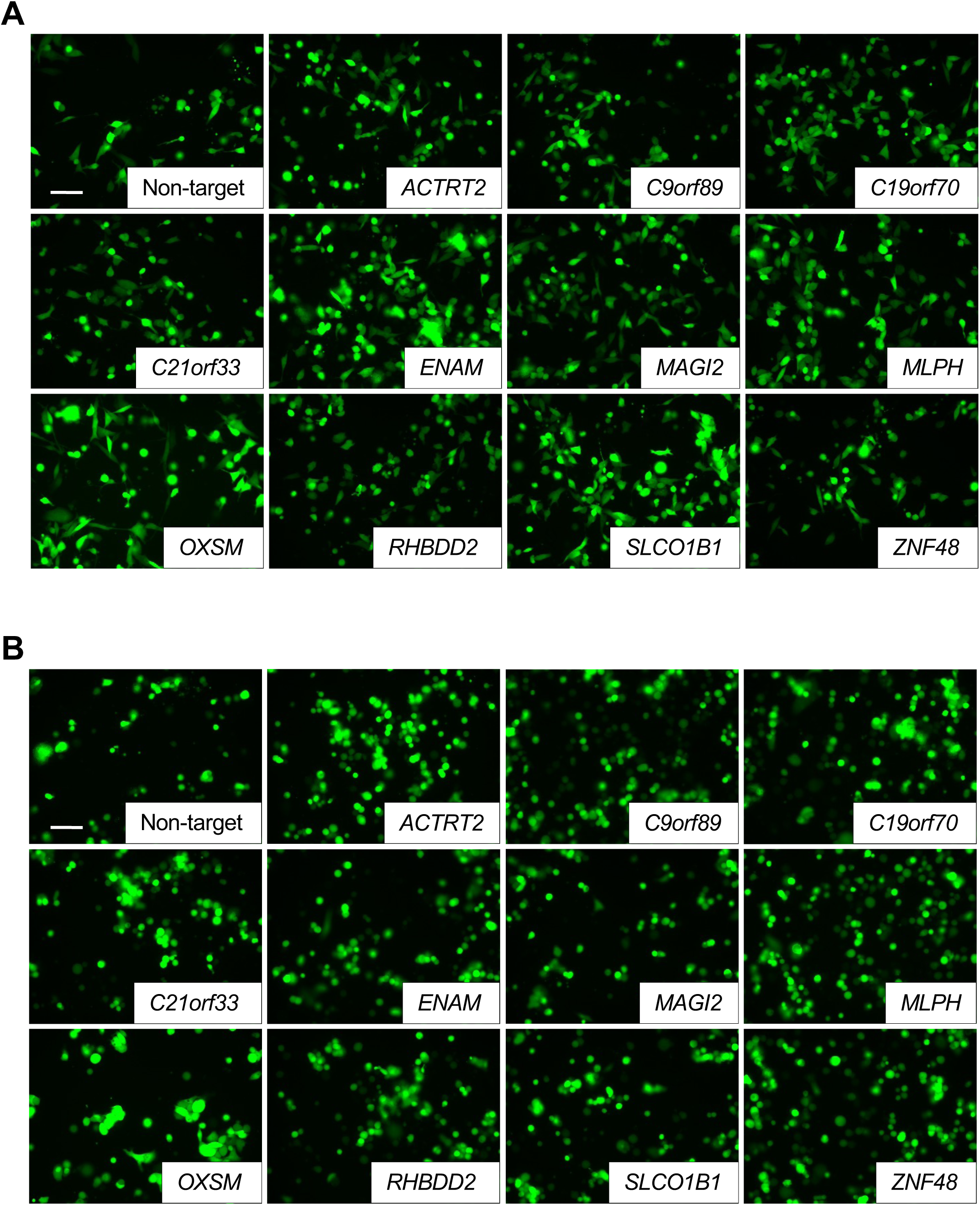
Representative images of GFP-positive cells in knockout mutant HEK293T-pancreatic cancer cells (MIA PaCa-2 (A) and SUIT-2 (B)) co-culture. (MIA PaCa-2 (A) and SUIT-2 (B)). After 168 h of exposure to gemcitabine (10 μM (A) or 3 μM (B)), images were captured under a green laser with a 20x objective field of view. Scale bar, 50 μm.

**Figure 4—figure supplement 1.**
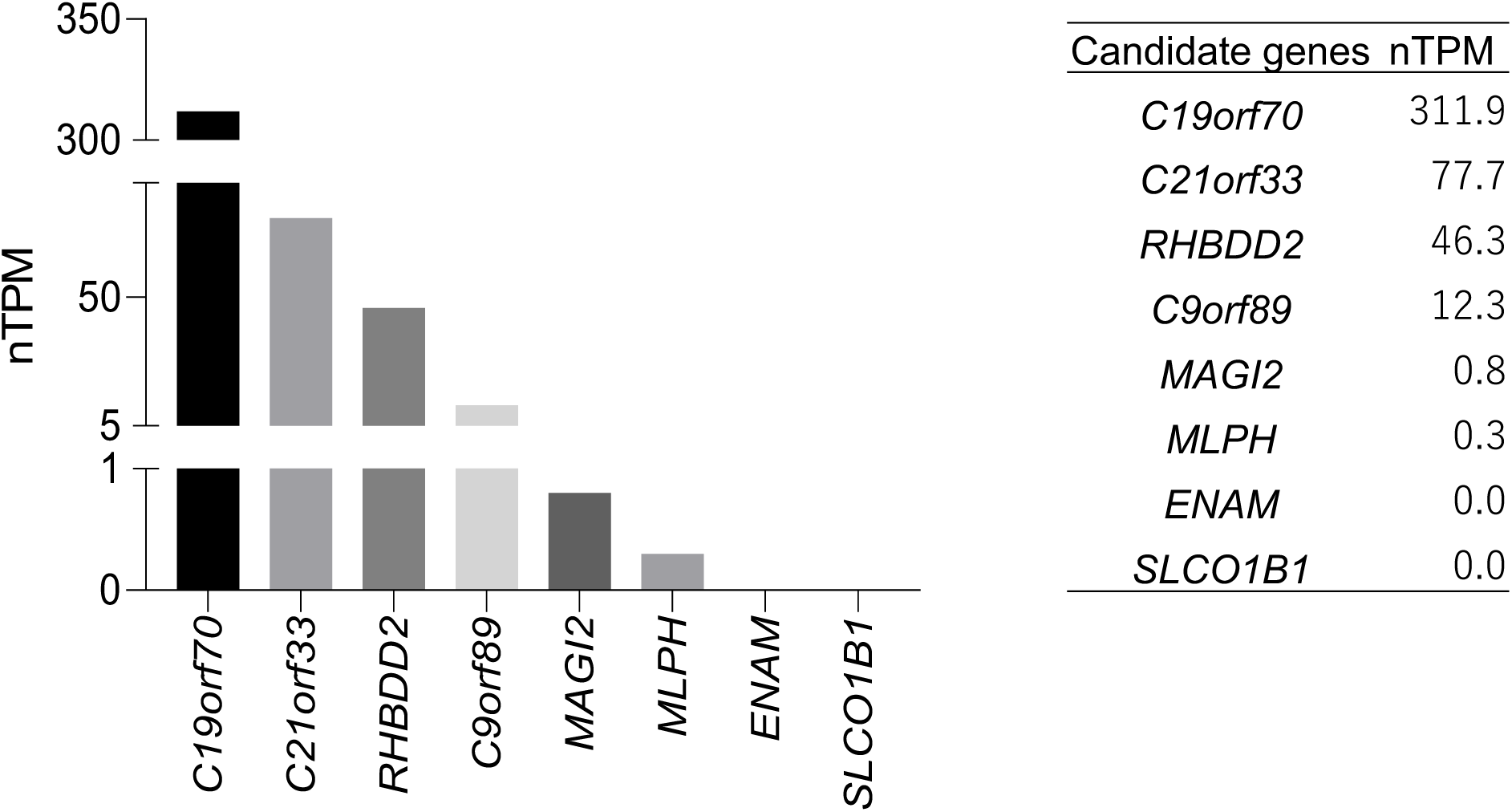
Transcriptional product expression of candidate genes in HEK293T. Transcriptional production (nTPM) ≥ 0.1 in HEK293T was defined as high expression based on transcriptional product expression data listed in the cell line RNA database of The Human Protein Atlas (https://www.proteinatlas.org/).

**Figure 4—figure supplement 2.**
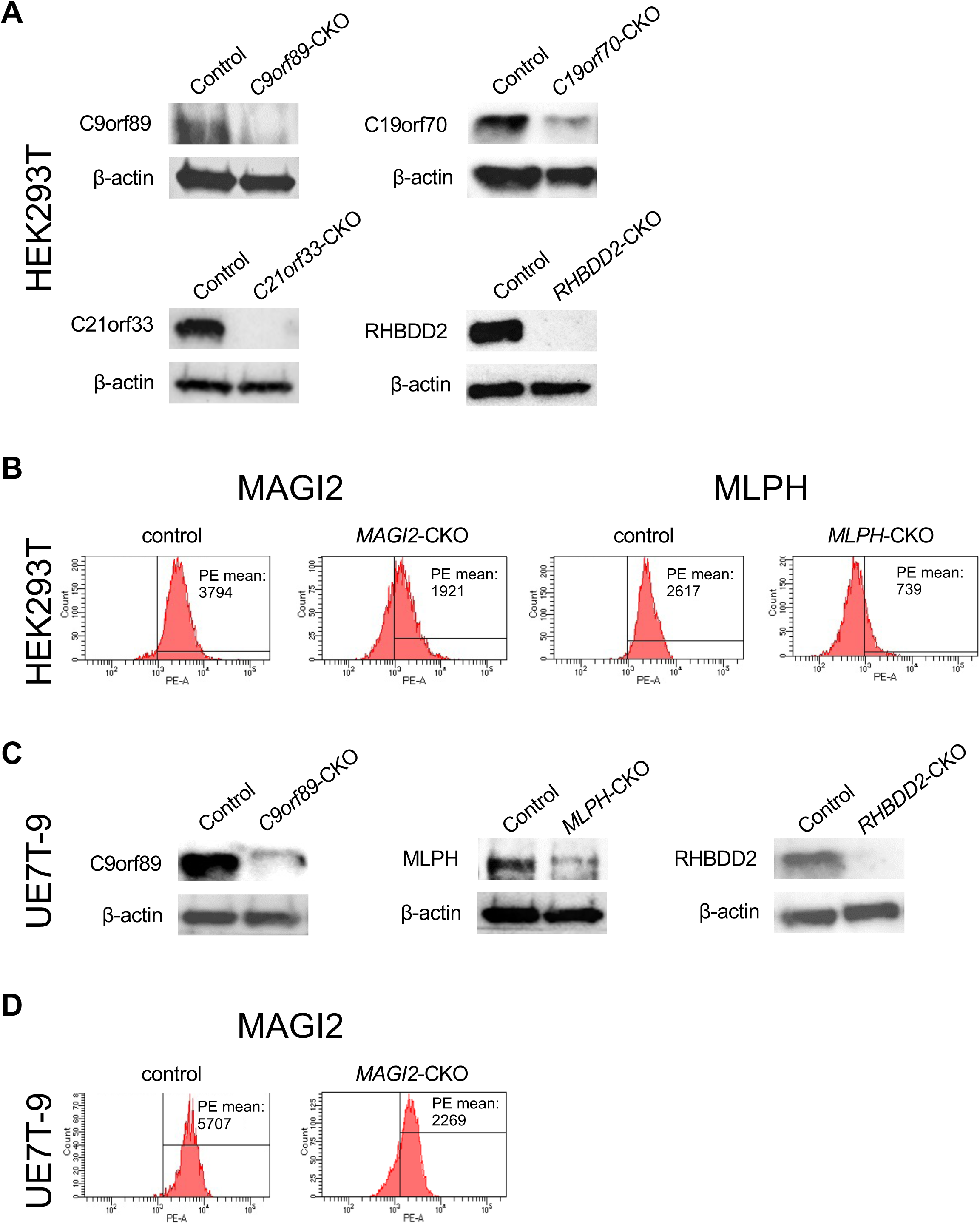
Validation of the loss of each candidate gene in HEK293T-CKO and UE7T-9-CKO clones. (A) Western blot analysis of C9orf89, C19orf70, C21orf33, and RHBDD2 of HEK293T-CKO clones. (B) Flow cytometric analysis of the expression of MAGI2 and MLPH in HEK293T-CKO clones. (C) Western blot analysis of C9orf89, MLPH, and RHBDD2 of UE7T-9-CKO clones. (D) Flow cytometric analysis of the expression of MAGI2 of the HEK293T-CKO clone.

**Figure 4—figure supplement 3.**
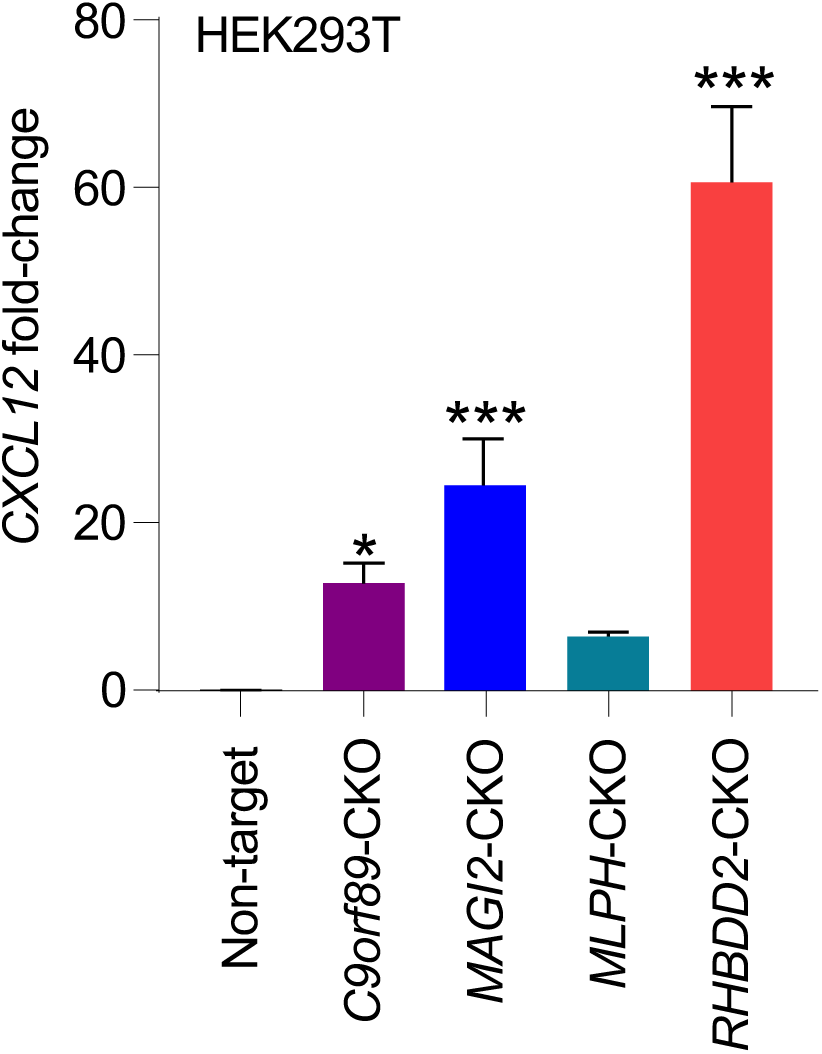
*CXCL12* expression in HEK293T-CKO clones. *CXCL12* expression was significantly up-regulated in HEK293T cells of *C9orf89*-CKO, *MAGI2*-CKO, and *RHBDD2*-CKO. The experiment was performed with biological triplication in the three independent experiments. Data are represented as mean ± SD. Statistical significance values were calculated by performing one-way ANOVA with Dunnett’s test. \**p* < 0.05 and \*\*\**p* < 0.001. Figure 4**—figure supplement 3—source data 1.** Values for the graph in Figure 4—figure supplement 3.

**Figure 4—figure supplement 4.**
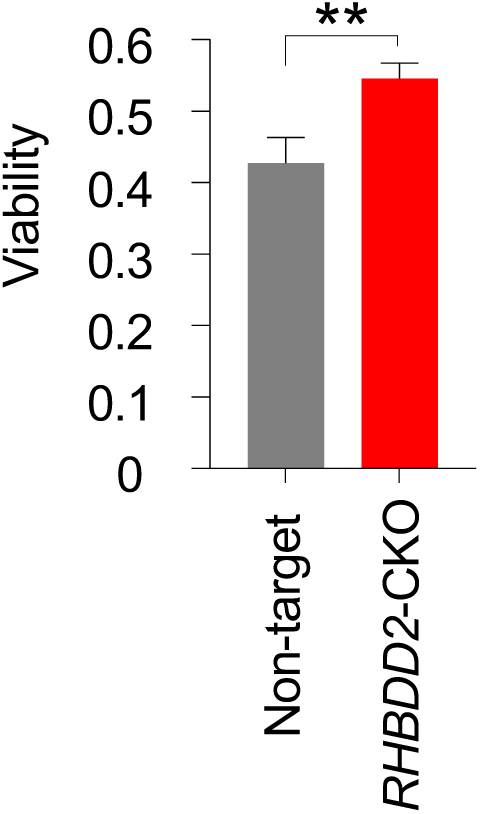
Supernatant of HEK293T *RHBDD2*-CKO induced cytarabine resistance in U937. U937 was cultured with the supernatant from HEK293T *RHBDD2*-CKO and treated with 250 nM of cytarabine for 48 h. The viability of U937 cultured with *RHBDD2*-CKO culture supernatant was slightly but significantly increased under cytarabine exposure. The experiment was performed with biological triplication in the three independent experiments. Data are represented as mean ± SD. Statistical significance values were calculated by performing two-tailed unpaired Student’s t-tests. \*\**p* < 0.01. Figure 4**—figure supplement 4—source data 1.** Values for the graph in Figure 4—figure supplement 4.

**Figure 5—figure supplement 1.**
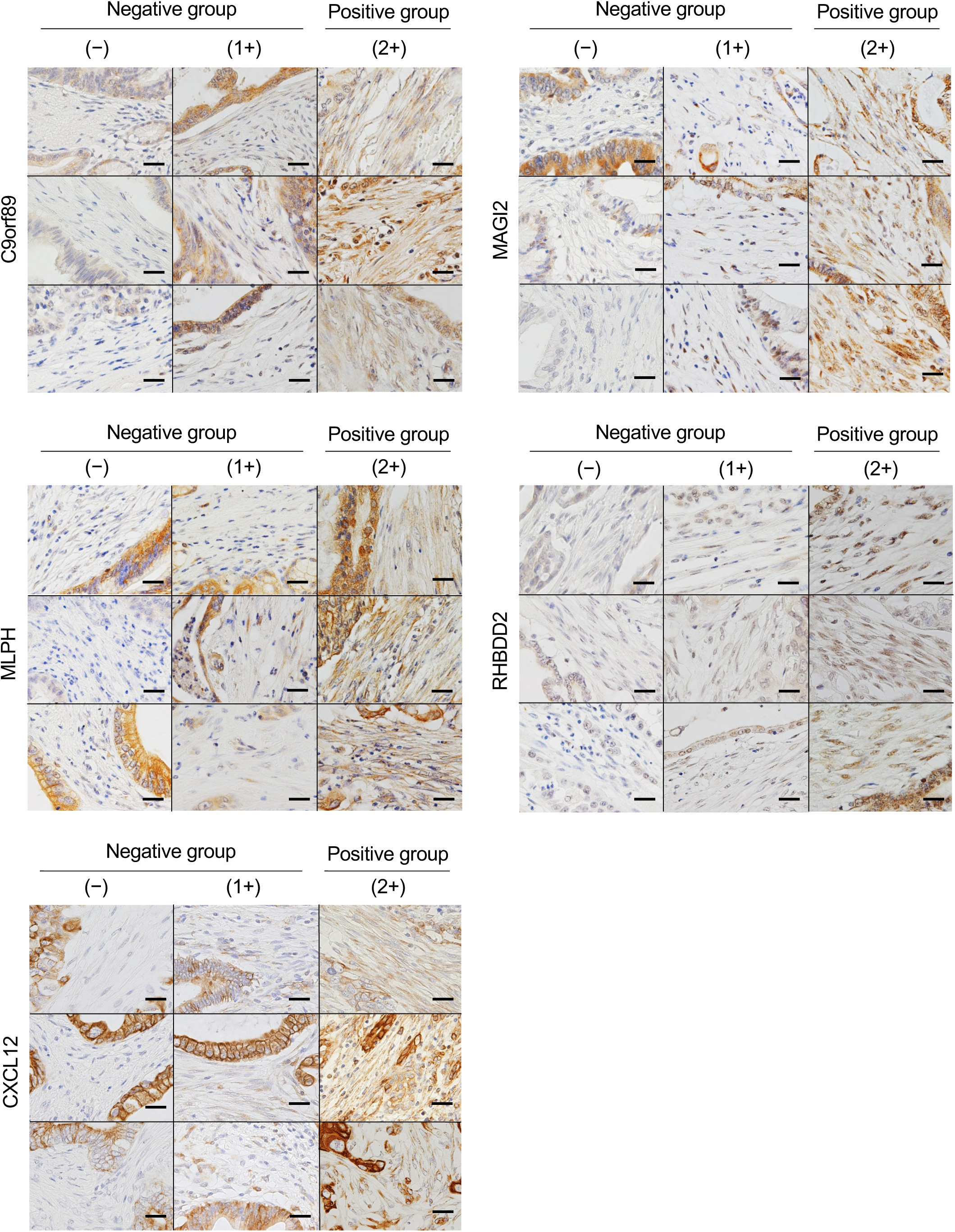
Immunostainability score of fibroblasts surrounding pancreatic carcinoma cells. The scoring of immunostainability was assessed based on the following criteria: not stained in fibroblasts was (-), positive in only a few fibroblasts or weakly positive were (1+), and strongly positive on most of the fibroblasts was (2+). Representative images of each antibody are shown. Based on scoring, (-) and (1+) were classified as negative groups, and (2+) was the positive group. Scale bar, 25 μm.

## Supplementary Files

**Supplementary File 1**

Up-regulated categories in HEK293T C9orf89-CKO in GSEA.

**Supplementary File 2**

Up-regulated categories in HEK293T MAGI2-CKO in GSEA.

**Supplementary File 3**

Up-regulated categories in HEK293T MLPH-CKO in GSEA.

**Supplementary File 4**

Up-regulated categories in HEK293T RHBDD2-CKO in GSEA.

**Supplementary File 5**

Universally up-regulated categories in HEK293T cells of C9orf89-CKO, MAGI2-CKO, MLPH-CKO, and RHBDD2-CKO.

**Supplementary File 6**

Summary of GSEA details of the category “CYTOKINE ACTIVITY” in HEK293T cells of C9orf89-CKO, MAGI2-CKO, MLPH-CKO, and RHBDD2-CKO.

**Supplementary File 7**

Summary of GSEA details of the category “CELL ADHESION MEDIATED BY INTEGRIN” in HEK293T cells of C9orf89-CKO, MAGI2-CKO, MLPH-CKO, and RHBDD2-CKO.

**Supplementary File 8**

Oligonucleotide sequence.

